# Somatic CRISPR editing of *Msh3* mitigates Huntington’s disease pathology in mice

**DOI:** 10.64898/2026.06.08.730940

**Authors:** Esaria Oliver, Marina Kovalenko, Mathilde Louçã, Andrew Jiang, Jordan Westerdahl, Kevin Correia, Benjamin Jones, Faaiza Saif, Nicole Romano, Ashna Sidhu, Tammy Gillis, Emanuela Elezi, Ryan Murtha, Ricardo Mouro Pinto, Vanessa C. Wheeler

**Affiliations:** Molecular Neurogenetics Unit, Center for Genomic Medicine, Massachusetts General Hospital, Boston, MA, USA; Department of Human Genetics, David Geffen School of Medicine, University of California, Los Angeles, Los Angeles, CA, USA (current address); Department of Neurology, Massachusetts General Hospital and Harvard Medical School, Boston, MA, USA; Medical and Population Genetics Program, the Broad Institute of M.I.T. and Harvard, Cambridge, MA, USA

## Abstract

Huntington’s disease (HD) is a fatal, dominantly inherited neurodegenerative disorder caused by a CAG repeat expansion in Huntingtin (*HTT*) exon 1. Further progressive CAG repeat expansion occurs in somatic cells, particularly in neurons, and drives the timing of clinical onset. Therefore, therapeutic strategies to slow somatic expansion are predicted to be disease-modifying. Somatic CAG expansion is driven by mismatch repair protein MSH3, a leading therapeutic target supported by human genetic data. To gain insight into the impact of targeting MSH3 at different stages of the disease process we used somatic CRISPR-Cas9 editing to knock out *Msh3* in *Htt*^Q111^ mice at ages of 6, 16, 24 weeks exhibiting progressively increasing somatic expansion. Intervention at all three ages slowed striatal CAG expansion, reduced nuclear huntingtin pathology and suppressed transcriptional dysregulation, with earlier intervention having greater impact. *Msh3* knockout also suppressed the production of the exon 1 *Htt1a* transcript. The results of our study provide important preclinical information relevant to an MSH3 therapeutic in humans that would be expected to impact a subset of cells in the brain, provide insight into the influence of timing of intervention on therapeutic effectiveness and deepen our understanding of how targeting MSH3 could alter the trajectory of HD.

## INTRODUCTION

Huntington’s disease (HD) [MIM: 143100]) is a fatal, dominantly inherited neurodegenerative disorder characterized by motor, cognitive and behavioral signs and symptoms, with selective vulnerability of medium spiny neurons (MSNs) in the striatum and projection neurons in the cortex ^1–3^. HD is caused by inheriting an expanded CAG repeat in *HTT* exon 1 whose length inversely correlates with age at onset (AAO) ^4^. The CAG repeat continues to expand in somatic cells, exhibiting high levels of expansion in the brain, particularly in vulnerable neuronal populations, and correlating with AAO ^5–10^. Human genome-wide association studies (GWAS) have revealed that AAO and other clinical landmarks are influenced by DNA repair genes that modulate somatic repeat expansion ^11–18^. The GWAS have provided strong support for a two-step model of HD pathogenesis in which the inherited CAG must first expand over an individual’s lifetime to reach a threshold(s) length, whereupon it triggers a toxic process(es) leading to cell death ^19,20^. A recent model based on single cell data indicates that expansion occurs in MSNs starting with a slow phase over decades until ∼80 CAGs, followed by a rapid phase over years, until the repeat reaches ∼150 CAGs, at which point, MSNs exhibit cell-autonomous transcriptional dysregulation, lose their neuronal identity and degenerate within a few months ^5^. These observations raise the possibility that maintaining somatic CAG repeat length below a critical pathogenic threshold, through timely intervention, could prevent clinical disease manifestation in HD mutation carriers.

A compelling therapeutic target to slow somatic expansion is MSH3. MSH3 preferentially recognizes CAG loop-outs as part of the MutSβ (MSH2-MSH3) heterodimer, initiating a DNA repair process that leads to expansion ^15,17,21–23^. Variants in *MSH3* that are associated with either increased *MSH3* expression or with decreased *MSH3* expression or loss of MSH3 function are associated with earlier or later clinical landmarks respectively ^11,13,24^, providing human-validated evidence that therapeutic MSH3 suppression will be disease-modifying. The low oncogenic liability of MSH3 ^25^ adds to its significance as a lead target.

In HD mouse models, constitutional homozygous *Msh3* knockout (KO) eliminates age-dependent somatic expansion, while heterozygous KO suppresses expansion ^15,26,27^. In HD CAG knock-in (KI) mouse models (*Htt*^Q111^ and *Htt*^Q140^), harboring inherited repeat lengths up to ∼140 CAGs, these genetic KOs, present from conception, slow the nuclear accumulation of mutant huntingtin protein (HTT) ^15,26^ and suppress transcriptional dysregulation in the striatum ^26^. In contrast, in a mouse model (*Htt*^Q175^) harboring ∼190 inherited CAGs, *Msh3* KO has no impact on these pathological and molecular phenotypes, indicating that a threshold CAG length has been exceeded beyond which stopping expansion has no phenotypic benefit ^27^. This observation is consistent with predictions from human single cell data ^5^.

Collectively, these data establish that the therapeutic benefit conferred by targeting somatic CAG expansion is shaped by two interdependent factors: the timing of intervention relative to disease progression, and the extent of repeat expansion already accumulated, findings with direct implications for the design of clinical trials in HD. In this study we have addressed the question of intervention timing in *Htt*^Q111^ mice in the context of an inherited CAG repeat of ∼120 CAGs that is rapidly expanding. To do this we employed a previously established *in vivo* somatic CRISPR-Cas9 editing system to KO *Msh3* in the brain using systemic delivery of PHP.eB adeno-associated virus (AAV)-mediated single guide RNAs (sgRNAs) delivery ^17^ at progressively older ages in order to compare the relative impacts on the repeat length-dependent suppression of somatic expansion and downstream pathological and molecular disease hallmarks. We demonstrate that *Msh3* KO suppresses striatal somatic CAG expansion and associated disease hallmarks, but that both the degree of expansion suppression and the extent of pathological rescue are greatest when intervention occurs earlier. These findings indicate an age-dependent therapeutic window for somatic expansion-targeting strategies and support the prioritization of early intervention in the design of HD clinical trials.

## RESULTS

### CRISPR-Cas9 knockout of *Msh3* at multiple ages suppresses MSH3 protein

Figure 1 outlines the experimental strategy. Heterozygous *Htt*^Q111/+^ mice or *Htt*^+/+^ littermates (referred to here as Q111 and WT, respectively), both constitutively expressing Cas9, were injected systemically at 6, 16 or 24 weeks of age with the PHP.eB adeno-associated virus (AAV9) variant ^28^ expressing a sgRNA targeting *Msh3,* or a control PHP.eB harboring empty vector. Cohorts of mice injected at 6 and 16 weeks of age were sacrificed at 24 weeks of age, and cohorts injected at all three timepoints were sacrificed at 40 weeks of age. Additional untreated cohorts were collected between 6 and 40 weeks of age. Tissues were processed for DNA-, RNA-, protein-and histology-based analyses of the striatum. Intervention time points were selected at which there are low (6 weeks), moderate (16 weeks) or high levels (24 weeks) of CAG expansion in the striatum ^17^, and analysis time points were selected for which molecular and histological phenotypes had been previously well characterized in this line ^15,16,29,30^.

**Figure 1:**
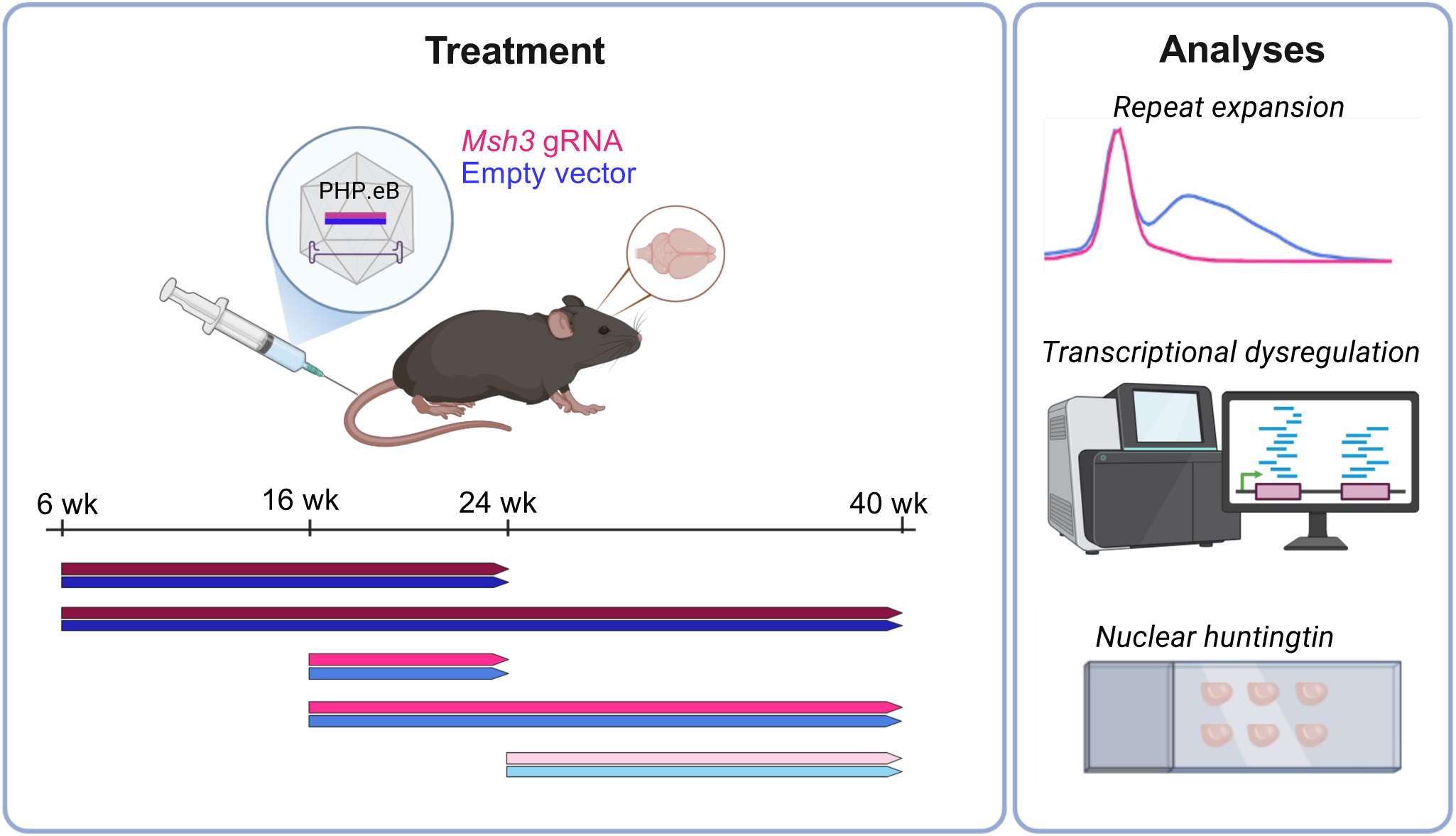
Outline of experimental timeline and analyses. Cas9-expressing *Htt*^Q111/+^ (Q111) and *Htt*^+/+^ (WT) mice were injected with PHP.eB-sgRNA targeting *Msh3* or with an empty-vector control at 6, 16, or 24 weeks of age and sacrificed at 24 or 40 weeks. Untreated cohorts were collected at matching timepoints. Phenotypic analyses were performed on striatal tissues.

The *Msh3* sgRNA significantly suppressed MSH3 protein levels in the striatum relative to empty vector injected mice in each treatment cohort (Fig. 2; 6-24 weeks: 79 % reduction, p < 0.0001; 16-24 weeks: 76 % reduction, p < 0.0001; 6-40 weeks: 89 % reduction, p = 0.0001; 16-40 weeks: 66 % reduction, p < 0.0001, 24-40 weeks: 63 % reduction, p < 0.001 [unpaired student’s t-tests]). The decreasing efficacy of MSH3 suppression with increasing intervention age approximately reflects a decrease in editing efficiency with age (Fig. S1A). This is likely due to a reduced viral dose: weight ratio in older mice as all animals received a fixed viral dose (1E+12 viral genomes). As previously described ^17^, and as shown again here (Fig. S1B,C), the impact on MSH3 protein level is high relative to the percentage of frameshift edits, suggesting an underestimation of inactivating mutations. Further, as also discussed previously ^17^, comparison of the impact of *Msh3* CRISPR KO with those of homozygous and heterozygous constitutional *Msh3* null mutations in the same mice ^15^ indicates that we are largely achieving biallelic inactivation.

**Figure 2:**
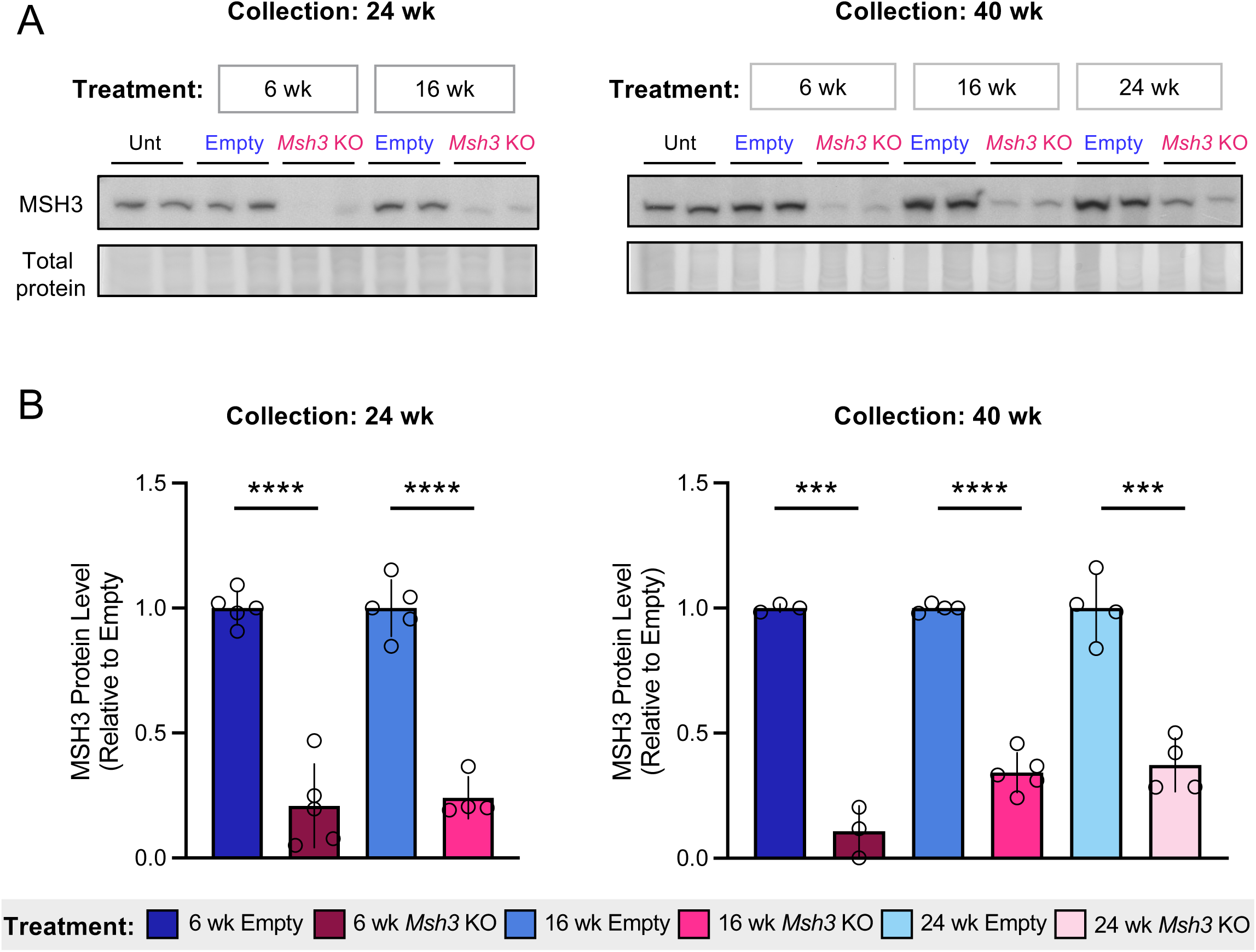
CRISPR-Cas9-mediated *Msh3* knockout reduces MSH3 protein in the striatum. A. Representative western blots showing MSH3 protein and total protein loading control in striatal lysates from Q111 mice injected with PHP.eB *Msh3* or empty vector control across treatment cohorts. Unt: untreated mice. Treatment ages are in grey boxes and analyses ages are shown above. B. Quantification of MSH3 protein levels normalized to total protein and expressed relative to the mean of the empty vector controls for each treatment group. Treatment groups are shown below the graphs and analyses ages are shown above. Unpaired Student’s t-tests: **** p<0.0001; *** p<0.001. N=3-5 Q111 mice per condition. Bars show mean +/-standard deviation.

### CRISPR-mediated *Msh3* knockout at multiple ages suppresses somatic expansion

We determined a somatic expansion index ^31,32^ from striatal genomic DNA of untreated, empty vector and *Msh3* sgRNA-treated mice, capturing a measure of somatic expansion in a bulk population of DNA molecules. In untreated and empty vector-treated mice, we observed an increase in expansion index at 40 weeks relative to 24 weeks as expected, and no effect on expansion in the empty vector-treated mice in comparison to untreated mice (Fig. 3A, Fig. S2). CRISPR-mediated *Msh3* KO significantly reduced somatic expansion index relative to empty vector-injected mice in each treatment cohort (Fig. 3B; 6-24 weeks: 58 % reduction, p < 0.0001; 16-24 weeks: 19 % reduction, p < 0.0001; 6-40 weeks: 84 % reduction, p < 0.0001; 16-40 weeks: 43 % reduction, p < 0.0001; 24-40 weeks: 18 % reduction, p < 0.0001 [unpaired student’s t-tests]), with expansion indices correlating with MSH3 protein level (Fig. 3C). For each analysis age (24 or 40 weeks), the relative reduction in expansion index with *Msh3* CRISPR KO compared to empty vector-treated animals decreased with progressively later ages of intervention. This is not simply explained by reduced *Msh3* editing at older intervention ages: Firstly, the variation in MSH3 protein reduction across the *Msh3* editing range, averaged for each cohort, is much more modest (reduced by 63-89 %) than the variation of impact on somatic expansion (reduced by 18-84 %) (Fig. S1B,C). *i.e.* relatively similar levels of MSH3 suppression can have different effects on somatic expansion index that depend on the intervention age. Secondly, while the average % *Msh3* editing is reduced with increasing age of intervention, analyses of expansion index as a function of % *Msh3* editing in individual mice reveals that each intervention timepoint impacts the expansion index differently across the same *Msh3* editing range (Fig. S1D).

**Figure 3:**
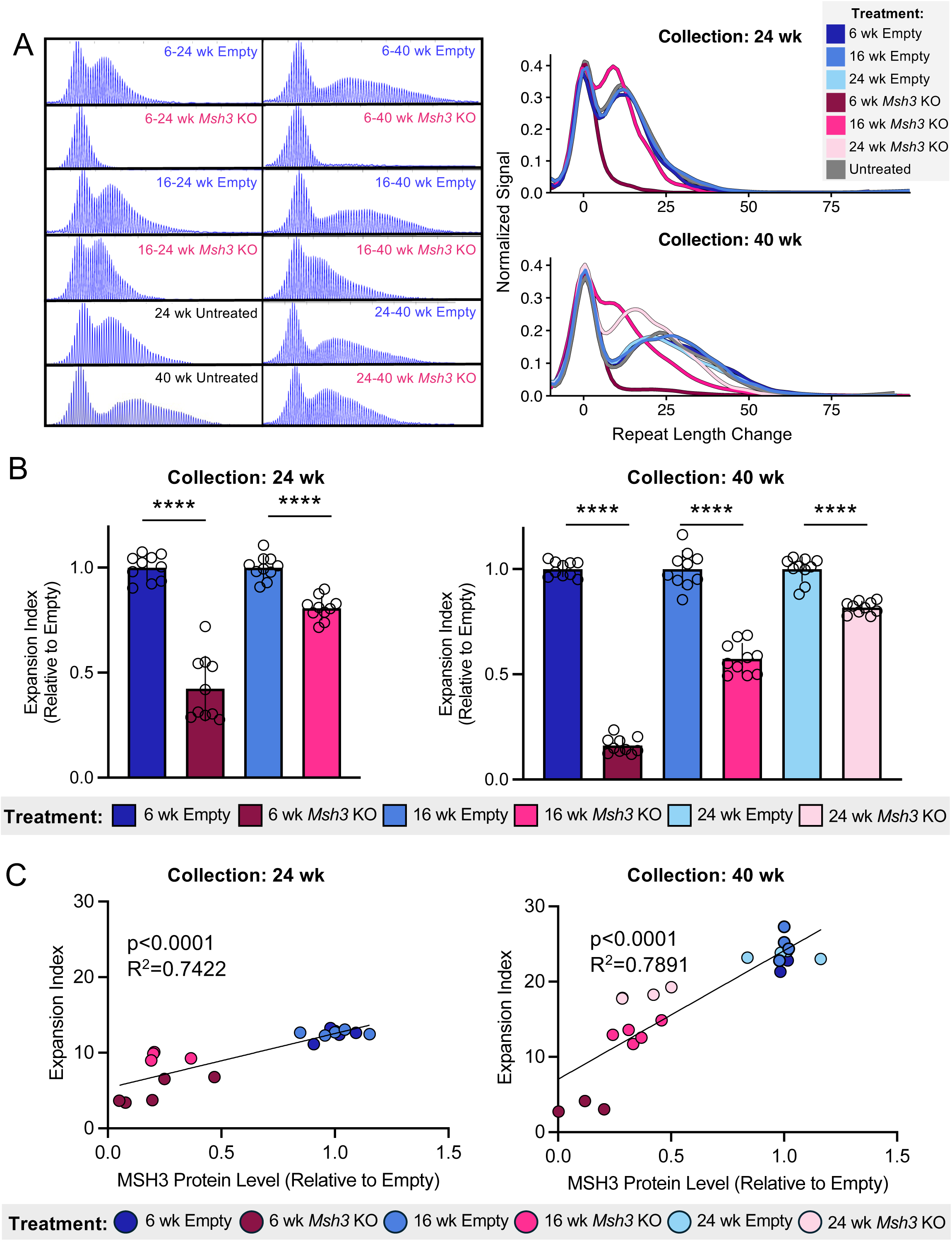
Effect of *Msh3* knockout at different ages on suppression of striatal CAG expansion. **A**. Left: Representative capillary electrophoresis traces showing CAG repeat length distributions in striatal DNA in *Msh3* KO, empty vector treated and untreated mice. Right: Overlaid traces normalized to the modal allele illustrate the shift in repeat length change between *Msh3* KO and empty injected or untreated mice. **B**. Quantification of somatic CAG expansion indices expressed relative to the mean of the empty vector controls for each treatment group. Treatment groups are shown below the graphs and analyses ages are shown above. N=10 Q111 mice per group. **** p<0.0001 (unpaired Student’s t-tests). Bars show mean +/- standard deviation. **C.** Correlation between somatic CAG expansion index and MSH3 protein level across all cohorts. MSH3 protein levels were normalized to total protein and expressed relative to the mean of the empty vector controls for each treatment group. Treatment groups are shown below the graphs.

To examine further the intervention age-dependent effects on CAG expansion we mapped expansion indices in our treatment cohorts onto a broader age trajectory of expansion that includes the 6- and 16-week intervention time points (Fig. 4A). From these data we estimate that the rate of change of expansion index, which is mathematically equivalent to the mean increase in CAG, is reduced from 0.63 CAGs/week in untreated mice to 0.037 CAGs/week (17-fold decrease), 0.27 CAGs/week (2.3-fold decrease) or 0.40 CAGs/week (1.6-fold decrease), in mice treated with the *Msh3* guide at 6, 16 or 24 weeks respectively. To account for pre-existing expansion accumulated prior to intervention, we also calculated the mean increase in expansion index from the age of intervention to the age of sacrifice for each cohort collected at 40 weeks (Fig. 4B). In empty vector-treated mice at 6, 16 or 24 weeks of age, the mean increase in expansion index was 20 CAGs, 17.1 CAGs and 10.5 CAGs. In comparison, *Msh3* KO at 6, 16 or 24 weeks of age resulted in a mean increase in expansion index of 1.5 CAGs, 7.1 CAGs and 6.3 CAGs, respectively. Thus, controlling for the expansion level at the time of intervention, these data demonstrate substantial suppression of new expansion regardless of intervention age.

**Figure 4:**
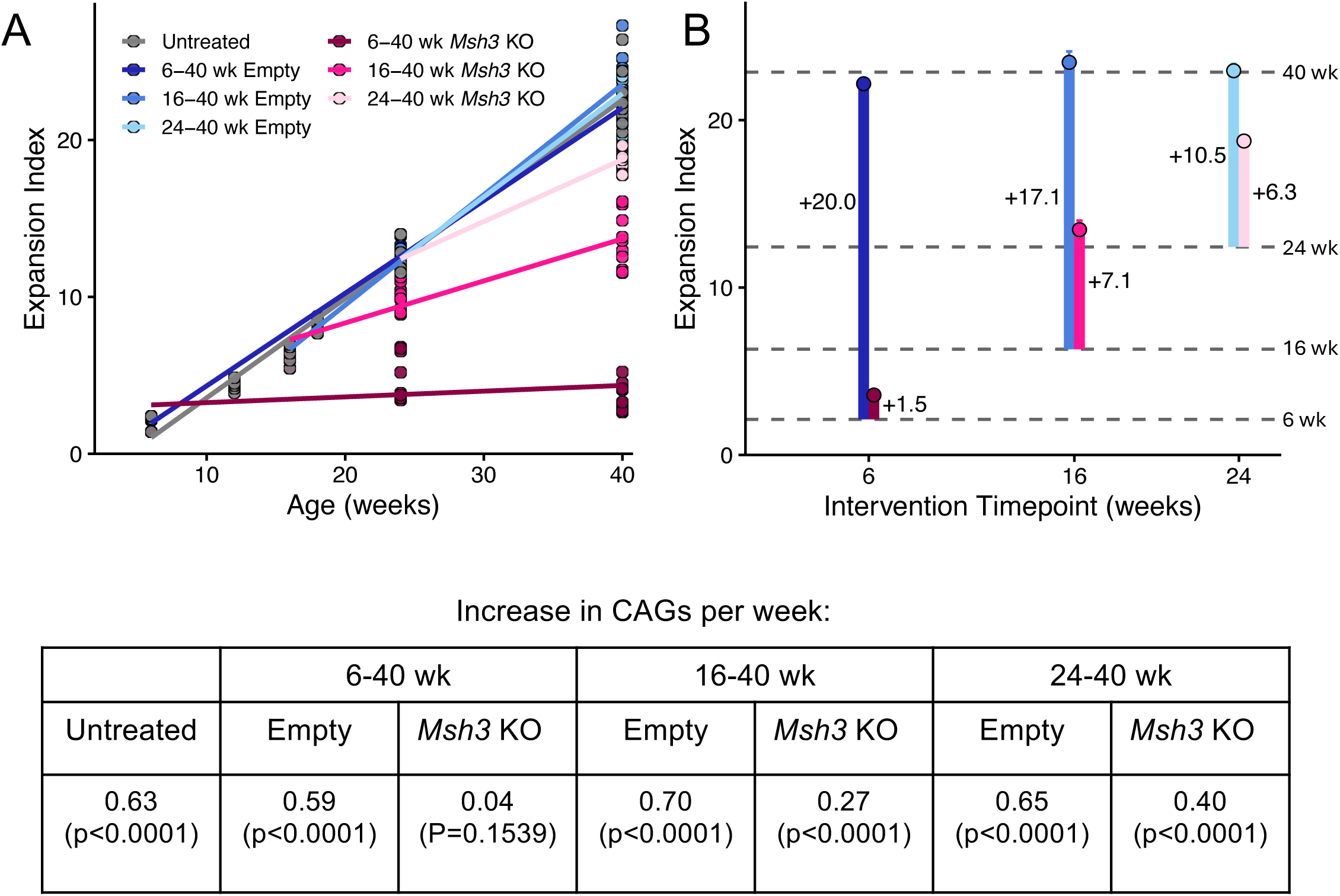
Age-dependent effects of *Msh3* knockout on the rate and magnitude of striatal CAG repeat expansion. **A.** Expansion indices from *Msh3* KO and empty vector-treated cohorts were mapped onto the age trajectory of somatic CAG expansion in untreated mice from 6 to 40 weeks of age. For injected mice, the starting points were anchored by expansion indices in untreated mice at 6, 16, and 40 weeks of age. In untreated Q111 mice, the rate of expansion was estimated at 0.63 CAGs/week. *Msh3* KO at 6, 16, or 24 weeks of age reduced this rate to 0.037 CAGs/week (17-fold reduction), 0.27 CAGs/week (2.3-fold reduction), and 0.40 CAGs/week (1.6-fold reduction), respectively. N=10 per condition for Q111 *Msh3* KO and empty vector and Q111 n=5-10 for untreated mice at each age. The table below shows the expansion rate (increase in CAGs per week) derived from the data in panel A. P-values show significance of deviation of the slope from zero **B**. The mean increase in expansion index from the time of injection to sacrifice at 40 weeks was calculated for each cohort. The vertical blue and pink lines show these increases for empty vector-treated and *Msh3* KO mice respectively. Horizontal dotted lines indicate expansion indices in untreated mice. N=10 per condition.

### Intervention age-dependent effect of CRISPR-mediated *Msh3* knockout on mutant huntingtin pathology

*Htt*^Q111^ mice display a characteristic age-dependent accumulation of mutant HTT species in the nuclei of striatal MSNs, first detected as a diffuse immunostain throughout the nucleus and subsequently as discrete intranuclear inclusions, detected by anti-HTT antibody mAb5374 (EM48) ^30,33,34^. We have previously shown that these histological phenotypes are CAG length-dependent ^33^^34, 35^and are suppressed by constitutional KO of *Msh3* and other MMR genes (*Msh2, Mlh1*) that drive somatic CAG expansion ^15,16,30^. By contrast, KO of genes that do not impact somatic CAG expansion (*Msh6, Xpc*) did not alter mutant HTT accumulation ^15^. These data indicate that these phenotypes are driven by, and provide a sensitive readout of, somatic CAG expansion in the striatum.

Therefore, we used mAb5374 immunohistochemistry to assess the impact of the *Msh3* CRISPR KO intervention on both total nuclear HTT intensity and intranuclear inclusions. At 24 weeks of age, mAb5374 staining is diffuse throughout the nucleus, while at 40 weeks of age, discrete inclusions are visible as well as diffuse stain (Fig. 5A, Fig. S3A). We used a protocol that is most sensitive to diffuse HTT to measure the total nuclear HTT intensity, which at 40 weeks includes the intensity contributed by both diffuse stain and inclusions (Fig. 5A). Diffuse mAb5374 staining intensity was quantified and to control for slide-to-slide variation, staining intensity in *Msh3* guide-treated mice was normalized to empty vector-treated mice on the same slide (Fig. 5B). Relative to empty vector, *Msh3* guide treatment at 6 weeks of age significantly suppressed nuclear HTT intensity measured at 24 weeks (44 % decrease, 2-tailed unpaired t-test, p = 0.0027), while *Msh3* guide treatment at 16 weeks resulted in a non-statistically significant trend towards reduced mAb5374 staining intensity. When measured at 40 weeks, *Msh3* guide treatment at all three time-points suppressed nuclear HTT intensity, though to progressively lesser extents with later intervention ages (Fig. 5B, treatment at 6 weeks: 77 % decrease, p < 0.0001; 16 weeks: 35 % decrease, p = 0.0045; 24 weeks: 14 % decrease, p = 0.03 [2-tailed unpaired t-tests]). We also applied a modified protocol to the 40-week mice that better detects the discrete intranuclear inclusions relative to the surrounding diffusely staining HTT (Fig. S3A). Inclusion number was also suppressed in *Msh3* guide-treated mice relative to empty vector-treated mice to an extent that depended on the intervention age (treatment at 6 weeks: 83 % decrease, p < 0.0001; 16 weeks: 46 % decrease, p = 0.0062; 24 weeks: 11 % decrease, p = 0.16 [2-tailed unpaired t-tests]) (Fig. S3B). Overall, the relative impacts on suppressing nuclear HTT phenotypes at the different intervention and analysis ages parallels the relative effects of the interventions on CAG expansion, consistent with the suppression of nuclear HTT being the consequence of somatic CAG expansion inhibition.

**Figure 5:**
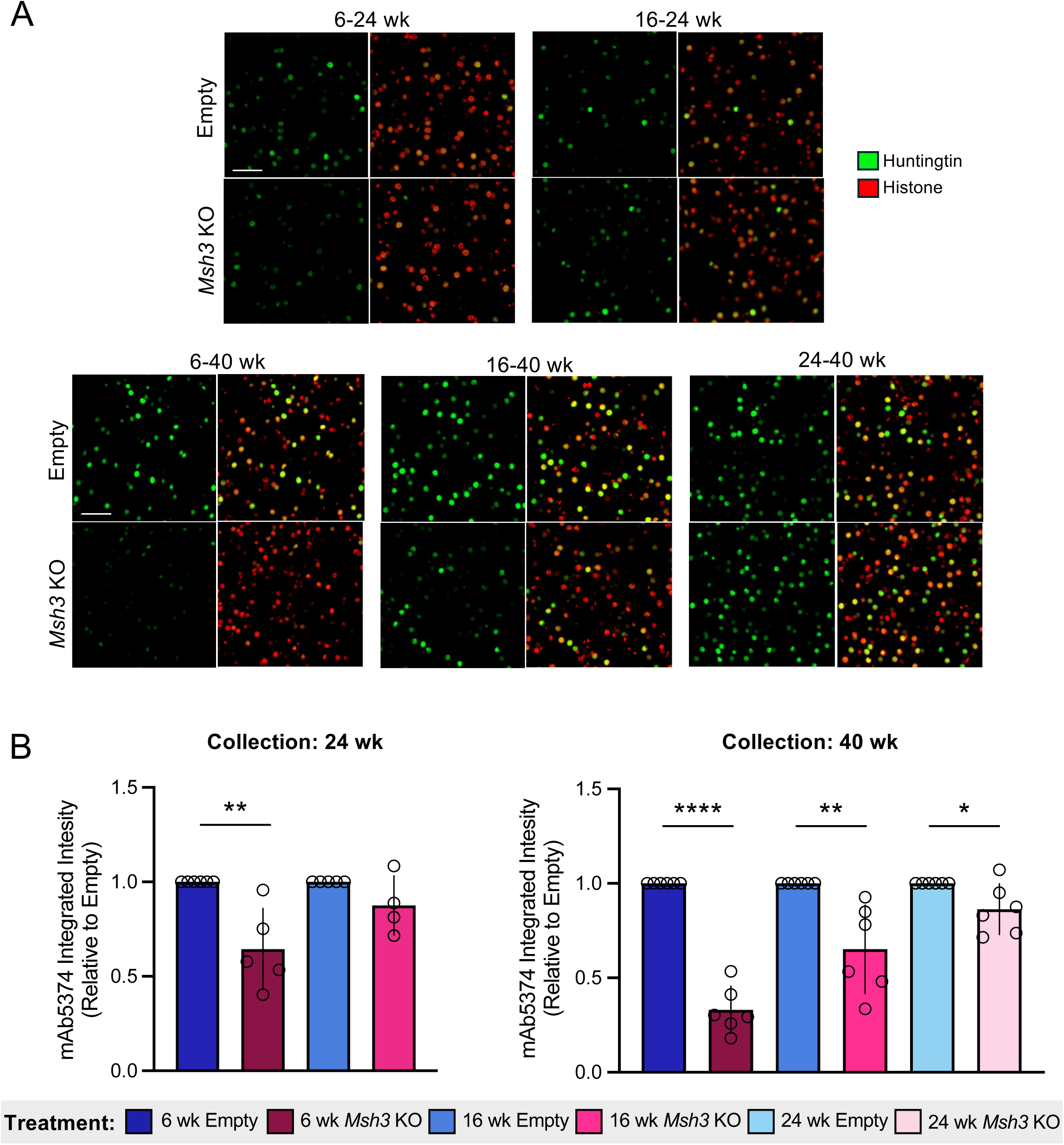
The effect of *Msh3* KO at different ages on nuclear huntingtin pathology. **A**. Representative images of Q111 striatal sections co-immunostained with anti-HTT mAb5374 (green) and anti-histone H3 (red; merged mAb5374 + histone H3 images are shown), illustrating diffuse nuclear HTT staining. Scale bar (white line) = 50 µm. **B**. Quantification of mAb5374 nuclear diffuse HTT staining intensity (see Methods) in *Msh3* KO and empty vector treated cohorts analyzed at 24 and 40 weeks of age. To control for slide-to-slide variation, staining intensity in *Msh3 KO* sections was normalized to the cohort-matched empty vector treated section on the same slide, and therefore all empty vector samples have a value of 1 (see Methods). In the 16-24-week group, a single outlier point with an mAb5374 integrated intensity value relative to empty of 1.9 that was driven by an anomalously low staining intensity of the empty vector-treated mouse on the matched slide was removed (outlier identification by ROUT method in GraphPad Prism v10, Q=10%). Unpaired Student’s t-tests: **** p<0.0001; *** p<0.001; * p<0.05. N=4-6 per condition. Bars show mean +/- standard deviation.

### CRISPR-mediated *Msh3* knockout at multiple ages partially rescues transcriptional dysregulation

Transcriptional dysregulation is a molecular hallmark in HD. Q111 mice exhibit transcriptional dysregulation ^29,36^, and across a series of HD KI mouse models, transcriptional dysregulation is CAG length-dependent ^36^, indicating that it can be modulated by factors that alter somatic repeat instability. Therefore, to investigate further the ability of the *Msh3* CRISPR KO to ameliorate disease expression we also performed RNA-sequencing (RNA-seq) of the striatum to analyze the impact on transcriptional dysregulation elicited by the Q111 allele. For each treatment group we performed RNA-seq in Q111 and WT striata in mice injected with the *Msh3* guide or empty vector, and in addition, in cohorts of 24-and 40-week Q111 and WT untreated mice. In untreated mice, we observed an age-dependent transcriptional dysregulation, with 1365 differentially expressed genes (DEGs) (FDR<0.1) in Q111 *vs*. WT striata at 24 weeks, increasing to 4598 DEGs at 40 weeks (Fig. S4). Similar numbers of DEGs were observed in Q111 *vs*. WT striata of mice that had been injected with the empty vector (Fig. 6A, Fig. S4A), indicating that the viral injection did not substantially alter the transcriptional response to the Q111 allele. However, comparisons of gene expression between empty vector-treated and untreated mice at each age revealed an intrinsic transcriptional response to the virus that was variable in effect (Fig. S4B, C). Therefore, in the subsequent analyses of the RNA-seq data we used the empty vector-injected mice as the control group, and henceforth, for simplicity, do not specifically refer to the empty vector.

**Figure 6:**
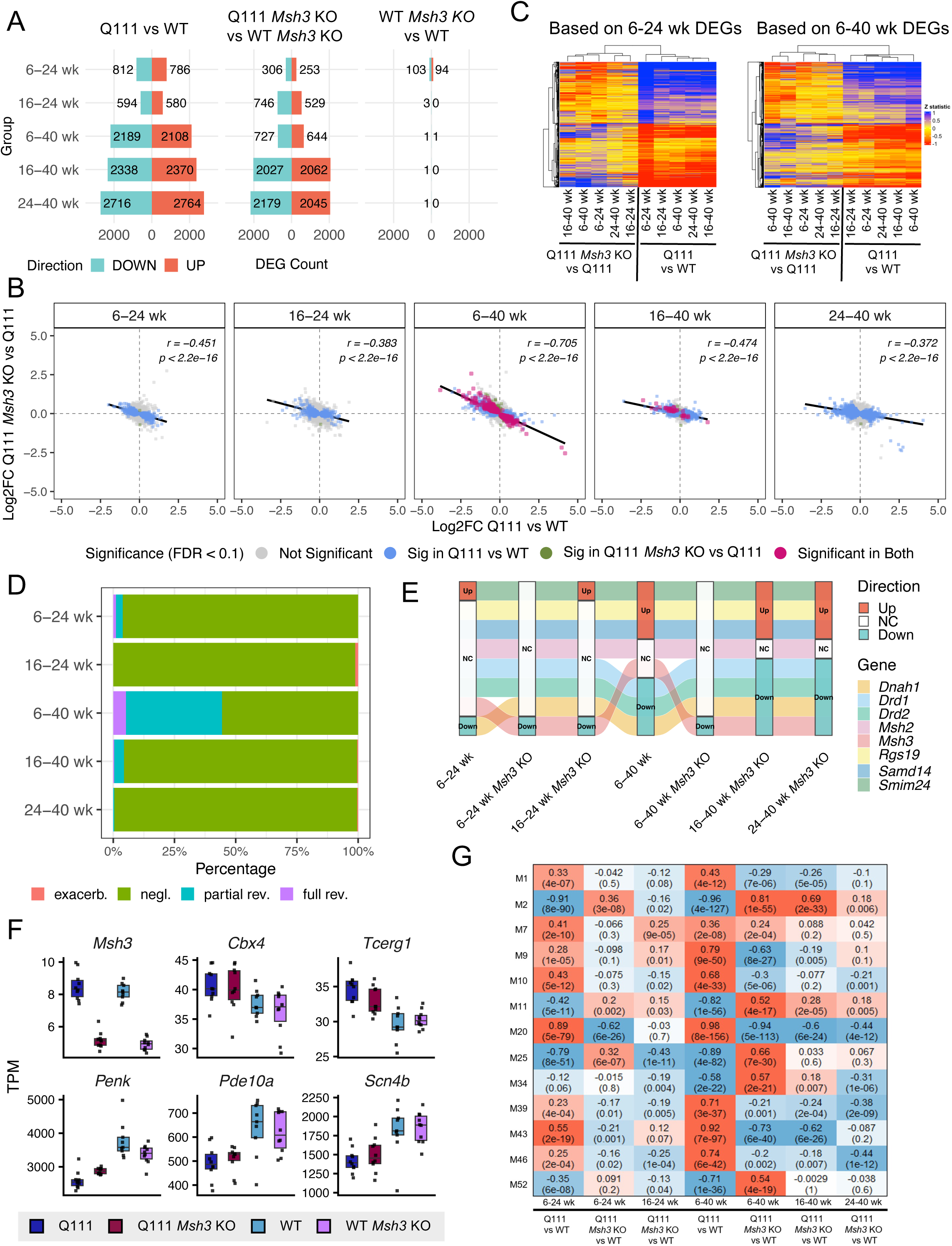
Partial rescue of transcriptional dysregulation. **A.** Numbers of up- or down-regulated differentially expressed genes (DEGs) (FDR<0.1) in the indicated comparisons. **B.** Gene expression changes (as log2 fold change [log2fc]) in Q111 *Msh3* KO *vs*. Q111 plotted against those in Q111 *vs*. WT for all groups (6-24 weeks, 16-24 weeks, 6-40 weeks, 16-40 weeks, 24-40 weeks). Key shows the number of DEGs significant in each comparison or in both. Genes in the bottom right and top left quadrants indicate Q111-dysregulated genes rescued by *Msh3* KO. **C.** Heatmaps of DEGs in Q111 vs WT and Q111 *Msh3* KO vs Q111 comparisons for mice analyzed at 24 weeks (left) or at 40 weeks (right). The DEGs represented in the left heat map are those defined by the 6-24-week Q111 vs WT comparison and the DEGs represented in the right heat map are those defined by the 6-40-week Q111 vs WT group. **D.** Reversal of gene expression using a posterior probability model based on the “STR266” signature. Color codes show exacerbation (excacerb.), negligible effect (negl.), partial reversal (partial rev.) or full reversal (full rev.). **E.** River plot for a subset of genes illustrating the relative change in gene expression by >20% transcripts per million (TPM) in Q111 (6-24 wk) and Q111 *Msh3* KO (6-24 wk, 16-24 wk) relative to 6-24 wk WT mice and in Q111 (6-40 wk) and Q111 *Msh3* KO (6-40 wk, 16-40 wk, 24-40 wk) relative to 6-40-wk WT mice. Based on the change in TMP, genes are classified as upregulated (up), downregulated (down) or not changed (NC). **F.** Gene expression values as TMP are plotted for *Msh3*, a subset of the up-and down-regulated “full” or “partial” reversal genes defined by the posterior probability model (*Cbx4, Penk, Pde10a, Snc4b*) and an HD modifier gene (*Tcerg1*) defined by the posterior probability, in the 6-40-wk group. **G.** Heatmap of weighted gene co-expression network analysis (WGCNA) based on striatal CAG length-dependent modules defined by Langfelder et al. ^36^ in Q111 vs WT comparisons and in Q111 *Msh3* KO vs Q111 comparisons for each intervention and analysis age. Red and blue indicate up- and down-regulated modules with p values in parentheses. Comparisons of *Msh3* guide-treated Q111 or WT are to empty vector-treated mice.

In comparison to the number of DEGs in Q111 *vs*. WT striata, the number of DEGs in Q111 *vs*. WT striata of *Msh3* guide-treated mice was reduced (Fig. 6A, compare left and middle panels), indicating that *Msh3* KO suppresses the response to the Q111 allele. In contrast to the strong transcriptional dysregulation elicited by the Q111 allele, *Msh3* KO in WT mice resulted in very few DEGs (Fig. 6A, compare left and right panels), indicating a low background effect of this genetic manipulation.

We further applied several analyses to assess the ability of *Msh3* KO to alter the transcriptional dysregulation in Q111 striatum. To evaluate the extent to which *Msh3* KO was able to normalize gene expression to levels in WT striata we plotted the gene expression changes (as log2 fold change [log2fc]) in Q111 *Msh3* KO *vs*. Q111 against those in Q111 *vs*. WT for all groups (6-24 weeks, 16-24 weeks, 6-40 weeks, 16-40 weeks, 24-40 weeks) (Fig. 6B). Points in the lower right and upper left quadrants show genes whose expression is changed in opposite directions in Q111 *Msh3* KO *vs*. Q111 and in Q111 *vs*. WT, indicative of the ability of the *Msh3* KO to revert or suppress the Q111-mediated gene expression changes. These plots revealed significant negative correlations for all groups, indicating an overall reversal of Q111-mediated gene expression changes by the *Msh3* KO. Earlier intervention ages and longer times between intervention and analysis produced stronger correlations and a greater number of DEGs for both the Q111 *vs*. WT and the Q111 *Msh3* KO *vs*. Q111 comparisons. Heatmaps based on the Q111 *vs*. WT DEGs in the 6-24-week group or in the 6-40-week group are shown in Figure 6C. For each, there were two distinct clusters that distinguished all Q111 vs. WT comparisons from all Q111 *Msh3* KO vs. Q111 comparisons, indicating consistent reversal effects of the *Msh3* KO across the different intervention ages. Quantifying previously described reversal correlation and reversal fraction statistics highlighted the strongest effect in the 6-40-week group with magnitudes comparable to those in constitutional heterozygous *Msh3* KO Q140 KI mice (Fig. S5) ^26^. We also assessed reversal of gene expression using a posterior probability model ^37^ based on the “STR266” signature comprising a set of commonly dysregulated genes across several HD models ^38^. Based on this model, only the 6-40-week group showed a substantial fraction of genes meeting criteria of “full” or “partial” reversal (Fig. 6D). Examples of trajectories of gene dysregulation in Q111 mice relative to WT mice across the different age and treatment groups for a subset of up- or down-regulated full or partial reversal STR266 signature genes (*Dnah1, Drd2, Smim24, Rgs19*) as well as *Drd1, Msh3 and Msh2* for comparison, are shown in Figure 6E. This indicates the relatively strong impact of the *Msh3* KO in the 6-40-week group. To visualize better the degree of rescue and the relative effects of *Msh3* KO in the Q111 background and WT background, gene expression values are plotted for the complete sets of full reversal genes in the 6-40, 16-40 and 24-40-week groups (Fig. S6), 6-40-week partial reversal genes (Fig. S7) and for HD modifier genes or candidates in the 6-40-week group (Fig. S8). A subset of these genes is shown in Figure 6F. These data highlight the impact of *Msh3* KO on moving gene expression in Q111 striatum to levels closer to those in WT striatum, and the lack of effect of *Msh3* KO on gene expression in WT mice. Gene expression of HD modifier genes or candidate genes is not strongly altered by Q111 or by *Msh3* KO (Fig. S8), though we find *Tcerg1* is upregulated by Q111, consistent with previous data ^36^ (https://www.hdinhd.org/) and that its upregulation is partially reversed by *Msh3* KO (Fig. 6F) This suggests a potential interaction between these two modifier genes that would be worth investigating in humans. Finally, to better capture global effects of the *Msh3* KO on gene expression beyond effect at the individual gene level, we performed Weighted Gene Co-expression Network Analyses (WGCNA) based on previously defined gene expression modules dysregulated in an HD knock-in CAG length allelic series ^36^ (Fig.6G, Fig. S9). CAG length-dependent modules (Fig.6G) were strongly dysregulated by Q111 at both 24 and 40 weeks of age. The direction of module dysregulation was strongly reversed for the majority of these modules in the 6-40-week *Msh3* KO group. While the reversal of module dysregulation was less at 24 weeks and eroded with later intervention ages, some modules (*e.g.* M1, M2, M10, M11, M20, M39, M46) were still significantly reversed by *Msh3* KO administered beyond 6 weeks of age. Amongst these, M2 (downregulated by Q111) is enriched in synaptic genes, GPCR and cAMP signaling genes and MSN identity genes and M20, M39 and M43 (upregulated by Q111) are enriched for protocadherin genes. Non-CAG length-dependent modules (Fig. S9) were less strongly dysregulated by Q111 and tended to be less reversed by *Msh3* KO, though strong reversal was seen for some (*e.g*. M16, M53).

Taken together, our analyses demonstrate that somatic KO of *Msh3* partially suppresses Q111-induced transcriptional dysregulation. Later intervention ages had reduced impact, consistent with the relative effects on suppression of somatic CAG expansion.

### CRISPR-mediated *Msh3* knockout suppresses *Htt1a*

Aberrant splicing of *HTT* exon 1 can occur due to increased usage of cryptic polyadenylation (polyA) sites in intron 1, resulting in a short polyadenylated mRNA (*HTT1a* or *Htt1a)* that is translated into an aggregation-prone exon 1 HTT protein (HTT1a) ^39,40^. As the aberrant splicing is CAG length-dependent, we tested whether *Htt1a* production is suppressed by *Msh3* KO. We used the RNA-seq data to quantify reads that would be contained in the *Htt1a* transcript based on alignment of RNA-seq reads from Q111 intron 1 (human sequence) to the cryptic polyA site in mouse intron 1 in Q111 mice (Fig. 7A) and performed a similar alignment to the equivalent region of the mouse genome in WT mice. We observed a substantial number of reads aligning to this region in Q111 mice, and very few in WT mice (Fig. 7B). This observation is consistent with these reads representing *Htt1a* transcripts ^39^, though we cannot rule out the possibility that they could also arise from other intron1-containing transcripts. However, given the agreement with published data ^39^ we assume that the intron1-to-polyA aligning reads largely represent *Htt1a* and refer to them henceforth as *Htt1a*. *Htt1a* expression was significantly suppressed by *Msh3* KO in 40-week Q111 mice (p < 0.001; 2-tailed unpaired t-test) and trended in the same direction in 24-week Q111 mice (Fig. 7B). As we had measures of somatic expansion, *Htt1a* and nuclear HTT from the same mice (and same hemi striatum for somatic expansion and *Htt1a*), we investigated relationships between them. To best capture striatal CAG lengths from our data, we calculated a mean striatal CAG length, which sums the modal CAG length in striatum and the expansion index (the latter being the mean increase in CAG length). *Htt1*a was significantly associated with mean striatal CAG length in both 24- and 40-week mice (Fig. 7C). Mean striatal CAG length explained 11.0 % of the variation in *Htt1*a in 24-week mice (p = 0.045) and 31.9 % of variation in *Htt1a* in 40-week mice (p=3.3E-6), with data points clustering according to the intervention timepoint (Fig. 7C). The number of nuclear HTT inclusions at 40 weeks (Fig. 7D) and the nuclear HTT intensity at 24 and 40 weeks (Fig.S10) were also significantly associated with mean striatal CAG length. Mean striatal CAG length accounted for 55 % of the variation in inclusion number (Fig. 7D) and 39-40 % of the variation in nuclear HTT intensity (Fig. S10), again with data points clustering according to the intervention timepoint. Finally, we also found positive associations between the nuclear HTT phenotypes and *Htt1a*, though these associations were weaker and only reached significance for the inclusion number in 40-week mice (Fig. 7E, Fig.S10).

**Figure 7:**
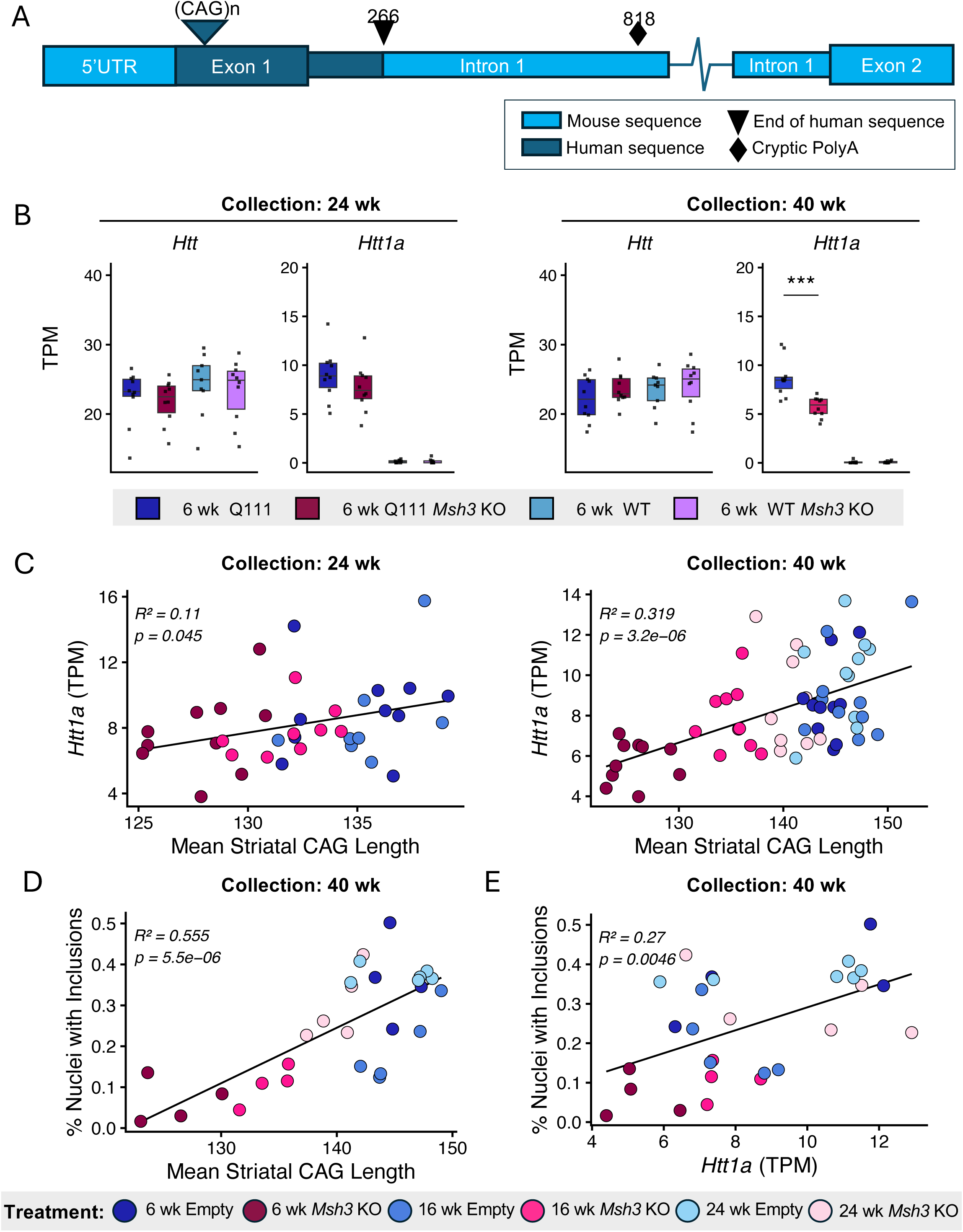
Rescue of *Htt1a* by *Msh3* CRISPR KO. A. Schematic of the genomic locus of the Q111 knock-in allele. Reads mapping from the beginning of intron 1 (position 1 in the numbering designation) to the cryptic polyadenylation site at position 818 were used to quantify *Htt1a* transcripts from the Q111 allele. Background intronic signal from the equivalent sequence in the WT allele in the Q111 heterozygotes would also be included, but is minimal, as seen in the WT mice. B. Box plots (median + 25th-75th percentile interquartile range) showing gene expression values as transcripts per million (TPM) for total *Htt*-aligned reads and for and *Htt1a* in the 6-24 and 6-40-week groups. N=9 or 10 per group. *** p<0.001 (unpaired Student’s t-test). C. Correlations between *Htt1a* and mean striatal CAG length across all groups. N= 8-10 per group. D. Correlation between % nuclei with mAb5374 inclusions and mean striatal CAG length in 40-week mice. N=4-6 per group. E. Correlation between % nuclei with mAb5374 inclusions and *Htt1a* in 40-week mice. N=4-6 per group. Comparisons of *Msh3* guide-treated Q111 or WT are to empty vector-treated mice.

Taken together, these data are consistent with the CAG-length dependent generation of *Htt*1a as the substrate for the aggregation-prone HTT1a protein in the nucleus and support the suppression of both *Htt1a* and nuclear mutant HTT being the consequence of somatic CAG expansion inhibition by KO of *Msh3*.

## DISCUSSION

Somatic expansion of the inherited *HTT* CAG repeat drives the onset of HD ^13^. Therefore, slowing or stopping somatic expansion is predicted to modify the course of the disease prior to clinical manifestation. *MSH3*, a genetic modifier of HD clinical phenotypes and key driver of somatic repeat expansion is a primary target that is being actively pursued for therapeutic benefit ^41^. Here, we have used somatic CRISPR-Cas9 editing in Q111 mice to gain insight into the potential to modulate repeat instability as well as downstream molecular and cellular pathogenic disease hallmarks when *Msh3* is knocked out in the striatum at different ages at which the mice exhibit progressively greater levels of somatic expansion.

We previously showed that constitutional homozygous genetic *Msh3* KO in Q111 mice stopped expansion ^15^, an effect that was recapitulated by somatic CRISPR KO at 6 weeks of age ^17^ and this study). Here, we have extended these observations to show that CRISPR KO at later ages (16 and 24 weeks) also significantly suppresses striatal CAG expansions. The impact of *Msh3* inactivation on the expansion index decreased with the later age of intervention age and was also dependent on the length of time after intervention. *e.g.* intervention at 6 weeks had a greater impact on the expansion index measured at 40 weeks than on the expansion index measured at 24 weeks.

An intervention age effect was also seen when controlling for the amount of somatic expansion at the time of intervention (Fig. 4B). It is unlikely that the reduced expansion index observed with progressively later intervention reflects a reduced ability of *Msh3* KO to suppress expansion of progressively longer repeats, as constitutional *Msh3* KO in HD knock-in mice harboring longer inherited repeats (*Htt*^Q140^ [Q140; ∼140 CAGs] ^26^ and z*Htt*^Q175^ [zQ175; ∼185 CAGs] ^27^ also stopped somatic expansion. *i.e.* in the absence of *Msh3* it appears that *HTT* CAG repeats are unable to expand, regardless of their length. Rather, as the expansion index quantified from bulk striatal tissue captures repeat length averaged across both edited and unedited MSNs - the cell-type in Q111 striata harboring the longest expansions ^30^- the effect of both intervention age and time between intervention and measurement reflects the extent to which the average striatal repeat length of the edited and unedited cells diverges from that in unedited striatum. *i.e*. for a given analysis age, the average repeat length diverges more in striata of mice treated earlier, and for a given intervention age, it diverges more as the mice are aged.

Consistent with our previous observations in constitutional *Msh3* KO mice ^15^, somatic CRISPR KO of *Msh3* at 6 weeks of age suppressed the intensity of diffuse nuclear mutant HTT immunostain at 24 weeks of age. At 40 weeks, KO of *Msh3* suppressed total nuclear HTT intensity (reflecting both diffuse stain and discrete inclusions) as well as the number of intranuclear inclusions. These 40-week phenotypes were also suppressed by *Msh3* KO at 16 and 24 weeks, with effect size decreasing with age of intervention, commensurate with the relative impacts on somatic expansion. These data are consistent with somatic expansion of the CAG repeat driving the accumulation of nuclear mutant HTT, a pathogenic hallmark of HD, and importantly, demonstrate that this process can be slowed by somatic inhibition of *Msh3* at multiple ages. Interestingly, in Q175 mice with ∼185 CAGs, *Msh3* genotype had no impact on nuclear HTT aggregation phenotypes detected with S830 antibody, indicating that once a repeat threshold has been surpassed, repeat stabilization can no longer modify these phenotypes ^27^. In Q140 mice, a potential CAG length threshold needed to elicit intranuclear mutant HTT inclusions, detectable with PHP-1 and mAb5374 anti-HTT antibodies, was determined to be ∼150 CAGs ^26^, which, notably, aligns well with a repeat length threshold proposed to drive cell-autonomous transcriptional dysregulation in MSNs in HD mutation carriers ^5^. The Wang et al. study_did not examine diffuse nuclear mutant HTT that precedes discrete intranuclear inclusions ^26^. In the current study, we estimate (Fig. 7, Fig. S10) a mean striatal repeat length of ∼131-139 CAGs at 24 weeks of age in empty vector-treated animals, stabilized to ∼125-131 CAGs in *Msh3* CRISPR KO animals treated at 6 weeks of age, and a mean striatal repeat length of ∼141-151 CAGs at 40 weeks of age in empty vector-treated animals, stabilized to ∼123-130 CAGs in *Msh3* CRISPR KO animals treated at 6 weeks of age. Given that we do not observe complete rescue of diffuse nuclear mutant HTT at 24 weeks of age (Fig. 5), this suggests that the threshold for detection of diffuse mAb4374-positive nuclear HTT species is lower than the above ∼150-CAG threshold. While the mean striatal CAG length metric from bulk PCR amplicons is likely an underestimate, a threshold of <150 CAGs also supported by our previous data in Q111 constitutional *Msh3* KO mice ^15^. In that study, the inherited repeat length of the mice was ∼100 CAGs, with an estimated MSN repeat length of ∼110 CAGs at 5 months of age. This elicited diffuse mAb4374-positive nuclear HTT that was suppressed by repeat stabilization to ∼100 CAGs in *Msh3* null mice. Due to the use of different HTT antibodies and detection methods that may differ in their ability to detect specific nuclear HTT species it is difficult to compare directly across studies, but overall, the data indicate that the repeat needs to exceed at least 100 CAGs in MSNs to detect any nuclear HTT species.

Notably, we find evidence that *Msh3* KO can suppress the expression of *Htt1a*. This was most apparent in the 6-40-week group and trended in the same direction in the other groups. The approximate concordance between the relative impacts of *Msh3* KO on somatic expansion, nuclear HTT phenotypes and *Htt1a* expression across the different groups appears consistent with the hypothesis that somatic expansion drives *Htt1a* expression that provides a substrate for an aggregation-prone HTT1a exon1 protein^39^, though does not rule out the idea that nuclear mutant HTT species may also arise independently of *Htt1a* production.

We also show that CRISPR-mediated *Msh3* KO partially suppresses global transcriptional dysregulation elicited by the Q111 allele. For all combinations of intervention and analysis ages, there were significant inverse correlations between the direction of gene expression changes elicited by the Q111 allele and by the *Msh3* KO in Q111 mice (Fig. 6), consistent with gene expression being reverted by *Msh3* KO closer to levels in WT striatum. Again, in line with the relative impact on somatic expansion, the effect on transcriptional dysregulation depended on the intervention age and the interval between intervention and analysis. Thus, we only observed a substantial number of genes significantly altered by *Msh3* KO in Q111 striatum when mice were treated at 6 weeks and analyzed at 40 weeks. For this condition, 44% of the “266STR” signature genes detected met criteria for either full or partial reversal. An interesting comparison of our data is with those in a recent study in which Q111 mice were treated at three ages (2, 5 and 8 months) with a divalent siRNA targeting *Msh3* ^42^. Analyses of rescue of transcriptional dysregulation in 12-month striata using the same posterior probability method and significance threshold showed ∼22% of “266STR” signature genes detected were fully or partially reversed by the siRNA treatment. Overall, these data align well with those in our study, with the greater rescue with *Msh3* CRISPR administered at 6 weeks of age appearing to reflect a stronger impact on somatic expansion; expansion in the *Msh3* siRNA-treated mice qualitatively appears close to levels expected in 3-4-month mice, while expansion in the *Msh3* CRISPR-treated mice is close to levels in 6-week untreated mice. Differences in instability suppression likely relate to the age of earliest intervention, *Msh3* knockdown versus KO, and expansions that might occur over time between *Msh3* siRNA dosing.

Similar to the nuclear HTT phenotypes, and in line with observations in Q140 and Q175 mice ^26,27^, our data are consistent with somatic CAG expansion beyond a threshold length driving transcriptional dysregulation in the striatum, and the potential for *Msh3* inhibition to suppress these molecular changes before the repeat has exceeded a threshold length. Ultimately, single cell-based analyses will be needed to gain insight into the repeat length threshold(s) needed for cell-autonomous gene dysregulation in the MSNs as well as in other cell types in the brain.

Overall, we demonstrate that therapeutic targeting of *Msh3* has the potential to alter early molecular and cellular events elicited by the *HTT* CAG repeat expansion. Translating effects in HD mouse models such as Q111, harboring very long repeats from birth, to humans with adult-onset HD is challenging. However, it is plausible that these mice might emulate the recently described rapid phase of somatic CAG expansion that is predicted to occur once the repeat has expanded beyond ∼80 CAGs ^5^. In HD mutation carriers, striatal MSNs undergoing rapid CAG expansion would be present in stochastic manner and increase with age. Our data indicate that stopping further CAG expansion at this point would impede the production of nuclear HTT inclusions and gene dysregulation that are predicted to presage eventual neuronal demise. Our data also indicate that earlier therapeutic intervention, targeting cells with lower repeats lengths, will have the greatest benefit. Our genetic study provides a preclinical model that is most relevant to an *MSH3* CRISPR-based therapeutic in humans that would also be expected to impact a subset of cells in the brain. It will also more generally inform additional preclinical work in HD around MSH3-targeted therapies, and, given the fundamental role of MSH3 as a driver of somatic expansion of many different disease-associated and other repeats ^43–47^, inform future preclinical work in other repeat expansion disorders.

## METHODS

### Mouse lines and injection with AAV vectors

Mouse experiments were carried out following the recommendations in the Guide for the Care and Use of Laboratory Animals, NRC (2010). All animal procedures were carried out to minimize pain and discomfort and following approved IACUC protocols of the Massachusetts General Hospital (MGH) or The Jackson Laboratory (JAX). *Htt*^Q111^ mice on the C57BL/6J (B6J) genetic background ^48^ were crossed with Cas9 knock-in mice constitutively expressing Cas9 nuclease under the direction of the CAG promoter (JAX strain 026179; B6J) ^49^ Male and female *Htt*^Q111/+^ and *Htt*^+/+^ (WT) mice, both heterozygous for the Cas9 knock-in allele, were shipped to MGH one week prior to injection. At 6, 16, or 24 weeks of age mice were administered via tail vein injection with 200 µl of 1E+12 PHP.eB vector genomes (VG). Male and female mice were each randomized to receive either PHP.eB expressing the *Msh3* sgRNA or PHP.eB carrying an empty vector. For each of the five age-treatment cohorts (6-24 weeks, 16-24 weeks, 6-40 weeks, 16-40 weeks, 24-40 weeks), ten *Htt*^Q111/+^ (five male and five female) and ten *Htt*^+/+^ mice (five male and five female) were injected with PHP.eB-*Msh3* sgRNA and ten *Htt*^Q111/+^ (five male and five female) and ten *Htt*^+/+^ mice (five male and five female) were injected with empty vector PHP.eB. Mice were aged to either 24 or 40 weeks of age and sacrificed for tissue harvesting. Additional cohorts of 24- and 40-week untreated *Htt*^Q111/+^ and *Htt*^+/+^ Cas9 heterozygous mice were collected in the same timeframe (ten mice [five males, five females] per genotype per age). Previously collected data from untreated *Htt*^Q111/+^ mice at 6, 12, 16 and 18 weeks were also specifically used for repeat instability comparisons, the predominant subset of which was previously described ^17^. CAG repeat lengths in all *Htt*^Q111^ mice ranged from 112 to 119 as determined from tail at weaning at Laragen, Inc. Mice were re-genotyped at MGH, in striatal DNA and were found to have CAGs ranging from 119 to 126. The MGH genotyping uses a standardized PCR assay (below) that sizes the repeat based on standards of known repeat length, and therefore we used these values as the more accurate measure of the repeat length in each mouse. Mice were transcardially perfused with ∼200 ml of PBS, and the striatum dissected from the left hemisphere of the brain, frozen on dry ice and stored at -80°C. The right hemisphere was fixed in 10% formalin at 4 degrees Celsius for 24 hours followed by three 10-minute rinses in Milli-Q^®^ water. They were then stored in 70% ethanol with 0.85% NaCl until they were processed for paraffin embedding ^30^.

### AAV constructs and viral production

We used a previously described plasmid vector for packaging in AAV (pAAV) ^17^ expressing a *Msh3*-targeting sgRNA under the control of the human U6 promoter and an mCherry reporter under the control of the CAGGS promoter, a hybrid promoter composed of the CMV immediate-early enhancer, CBA promoter, and CBA intron 1/exon 1 ^50^ The *Msh3* sgRNA was previously described and validated ^17^. pAAVs (*Msh3* sgRNA or empty vector) were packaged into PHP.eB capsids at the UMass Chan Medical School Viral Vector Core.

### DNA, RNA and protein extraction

DNA, RNA, and protein were extracted from the left striatum using TRIzol™ reagent (Thermo Fisher Scientific). Extractions were performed according to the manufacturer’s instructions, but including an extra 70% ethanol wash for both RNA and DNA. RNA was stored at -80°C and DNA stored at -20°C. Protein pellets suspended in Protein Wash 1 were stored for later use at -20°C. Before use, the protocol was continued with the addition of 3 x 10 seconds sonification pulses at power level 3.5 (550 Sonic Dismembrator, Fisher Scientific). Protein was solubilized in 1% SDS and then sonicated again for 3 x 10 seconds and heated at 50°C for ten minutes to maximize the amount of protein retrieved. DNA and RNA were quantified with a NanoDrop spectrophotomer, RNA quality was determined using Agilent Technologies 4200 TapeStation and protein concentration was determined using the BCA assay (Thermo Fisher Scientific).

### Quantification of CRISPR/Cas9 editing efficiency

PCR amplicons surrounding the *Msh3* target site ^17^ were generated using Phusion High-Fidelity DNA Polymerase (Thermo Fisher Scientific), purified using the MiniElute PCR Purification kit (Qiagen) and subjected to deep 2 x 150 bp paired-end next generation sequencing (NGS) on the Illumina MiSeq platform (DNA core, Center for Computational and Integrative Biology, MGH). Genomic editing was determined in each sample from a minimum of 10,000 NGS reads using CRISPResso v2 ^51^ using batch command with default settings and the following parameters: amplicon_min_alignment_score: 90; quantification_window_center: −3; quantification_window_size: 5. Editing efficiency was summarized as percentages of frameshift (non-multiple of 3bp indels) and non-frameshift (multiple of 3bp indels and SNVs) mutations.

### CAG repeat expansion analysis

Somatic CAG instability analysis was performed using a human-specific PCR assay that amplifies the *HTT* CAG repeat from the knock-in allele ^16^. The forward primer was fluorescently labeled with 6-FAM (Thermo Fisher Scientific) and products were resolved using the ABI 3730xl DNA analyzer (Thermo Fisher Scientific) with GeneScan 500 LIZ as internal size standard (Thermo Fisher Scientific). GeneMapper v5 (Thermo Fisher Scientific) was used to generate CAG repeat size distribution traces. CAG expansion indices were calculated as previously described ^31^ using the Trace Shiny program ^32^ with a 5% relative peak height threshold cut-off and a window of +100 CAGs beyond the modal allele. Expansion indices were determined relative to the modal allele in each trace. A mean striatal CAG was calculated by adding the expansion index to the modal CAG length in striatum.

### Western blot analyses

Protein extracts (40 µg) were resolved on 4–12 % Bis-Tris polyacrylamide gels (NuPAGE, Life Technologies) and transferred to 0.45 mm nitrocellulose membranes (Thermo Fisher Scientific). The membranes were blocked with 5% dry non-fat milk in Tris-buffered saline with 0.1% Tween 20 (TBST) and incubated with MSH3 antibody (1:5; conditioned medium from hybridoma MSH3MAB2(2F11) obtained from DSHB) in 5% milk/TBST overnight at 4°C. Horseradish peroxidase-conjugated sheep anti-mouse (1:5000; NA931, Cytiva) was used as secondary antibody. Signals were visualized using the enhanced chemiluminescence (ECL) detection system (Thermo Fisher Scientific). Densitometric analysis of protein levels was performed using ImageJ (Version 1.54g; National Institutes of Health, USA) software. Following background subtraction, protein levels were normalized by total protein amount in the same lane as determined by Novex Reversible Membrane Protein Stain (Thermo Fisher Scientific) performed prior to blocking. Protein levels were normalized relative to empty vector controls of the same intervention and collection timepoint on the same blot.

### Immunohistochemistry

PBS-perfused, 10% formalin fixed mouse brains (right hemispheres) were processed, paraffin embedded and sectioned at 6 mm coronally. The sections were deparaffinized, rehydrated and subjected to heat-mediated epitope retrieval in sodium citrate buffer (pH 6.0), followed by quenching of endogenous peroxidase with 0.3 % H_2_O_2_/methanol for 30 min at room temperature (RT) and blocked in 3 % normal horse serum (NHS) in Tris-buffered saline (TBS) for 1 hour at room temperature. For detection of diffuse nuclear HTT, incubation with huntingtin antibody mAb5374 (1∶100; MAB5374, Millipore) and histone H3 antibody as nuclear marker (1:500; ab1791, Abcam) in 1% NHS/TBS was carried out overnight at 4°C. The next day, sections were incubated with biotinylated horse anti-mouse IgG (1:200; BA-2000, Vector Laboratories) in 1% NHS/TBS for 1 h at RT. Then, mAb5374 signal was amplified using Biotin XX Tyramide SuperBoost™ Kit, Streptavidin (B40931, Thermo Fisher Scientific) according to the manufacturer’s protocol and detected with Streptavidin-Alexa Fluor™ 647 (1:500; Thermo Fisher Scientific) in 1% NHS/TBS for 1 h at RT. Histone H3 staining was detected with donkey anti-rabbit-Alexa Fluor™ 488 (1:500; A21206, Life Technologies) in 1 % NHS/TBS for 1 hour at room temperature, added together with Streptavidin-Alexa Fluor™ 647. For detection of HTT inclusions, sections were incubated with mAb5374 (1∶200) and histone H3 ab1791 (1:500) in 1 % NHS/TBS overnight at 4°C, followed by amplification of mAb5374 signal with Alexa Fluor™ 647 Tyramide SuperBoost™ Kit, (B40916, Thermo Fisher Scientific), according to the manufacturer’s protocol. Histone H3 was detected with donkey anti-rabbit-Alexa Fluor™ 488 (1:500) in 1 % NHS/TBS, 1 h at RT, added after the completion of Tyramide-Alexa Fluor™ 647 step. Sections were coverslipped with DAPI Fluoromount-G^®^ (SouthernBiotech). For quantification, fluorescent microscopy was performed with CellVoyager CQ1 (Yokogawa) of whole striatal sections at 10x magnification. Images were acquired with the same exposure times. Images used in the figures were acquired with a Leica DMi8 epifluorescence microscope at 10x, with the same exposure times.

### Quantification of immunostaining

Brains were paraffin embedded with either four or six brains per paraffin block, such that each included an empty vector control mouse and an *Msh3* treated mouse for each age-treatment group, allowing for block normalization. Each slide contained two consecutive sections from a single block. Quantification was carried out using Cell Profiler 4.2.8 ^52^. The striatum was manually outlined based on morphology. For diffuse mAb5374 immunostaining, the total (integrated) intensity of mAb5374 staining was measured in all H3-positive nuclei) and normalized by the total number of nuclei in the striatum, as determined by the number of histone H3-positive nuclei. This generated the mean integrated intensity of mAb5374 staining per nucleus. This value was then averaged per mouse across the two consecutive sections on each slide. To control for slide-to-slide variation in staining intensity, the averaged mean integrated intensity of mAb5374 of each mouse was normalized to that of the corresponding empty vector control mouse on the same slide. For inclusions, the total number of mAb5374-positive inclusions in the striatum was normalized by the total number of H3-positive nuclei. This value (number of inclusions per nucleus) was averaged across the two consecutive sections per mouse and then normalized (as above) to the corresponding value in the empty vector control mouse on the same slide.

### RNA-sequencing

cDNA libraries from 240 striatal samples were prepared using the Illumina TruSeq RNA Library Prep Kit and sequenced on the NovaSeq platform (150 bp paired-end reads). Adapter-trimmed reads were aligned using STAR ^53^ with default parameters to a custom Q111 mouse reference genome (based on mm39) incorporating the humanized knock-in sequence ^54^. Gene-level read counts were generated using HTSeq ^55^. We applied filtering criteria as described in Wang et al ^26^, retaining genes that exhibited >= 0.5 counts per million (CPM) in at least 8 samples (least number of samples in the smallest group) and identifying outliers as described ^26^. Nine outliers were removed, retaining N=10 per condition, with the exception of the following: 6-24wks WT Empty (n=9), 6-40wks WT Empty (n=9), 16-24wks Q111 Empty (n=8), 16-24wks Q111 Msh3 KO (n=9), 16-40wks WT Empty (n=8), 24-40wks Q111 Msh3 KO (n=9), and 40wks untreated Q111 (n=9). For comparison and visualization and of gene expression, expression levels were normalized to transcripts per million (TPM) to account for differences in both gene length and sequencing depth, allowing for direct comparison of transcript abundance across samples and genes or gene regions. Raw counts were divided by gene or gene region length (for *Htt1a*) in kilobases to calculate reads per kilobase (RPK). The sum of RPK values in each sample was then divided by one million to obtain a sample-specific scaling factor, and each gene’s RPK was divided by this factor to yield TPM values, ensuring comparability across samples.

Differential expression analysis was performed using DESeq2 1.48.2 with default arguments ^56^ in a model that included sex, RNA Integrity Number (RIN) and the first surrogate variable (SV) representing unaccounted for variation (surrogate variable analysis (SVA), R package sva ^57^ (as covariates. Differentially expressed genes between groups based on false discovery rate (FDR) <0.1 adjustment were assigned as significant.

### Analyses of *Msh3* knockout-mediated rescue of transcriptional dysregulation

We applied multiple statistical approaches to quantify the extent of transcriptomic rescue induced by *Msh3* KO. For each condition, we compared the numbers of significantly differentially expressed genes (DEGs) in Q111 (empty vector) versus WT (empty vector) with those in Q111 *Msh3* KO versus Q111 empty vector. Genes exhibiting reversal showed opposite directions of fold-change between the two contrasts, whereas exacerbated genes showed concordant fold-change directions. We employed a Bayesian statistical framework to estimate, on a gene-by-gene basis, the posterior probability that *Msh3* KO reverses disease-associated transcriptional dysregulation. This approach was adapted from Marchionini et al. ^37^ and has been applied to assess *Msh3*-mediated transcriptional rescue in the zQ175 and Q111 models ^27,42^. Briefly, this approach models the effect of *Msh3* KO as a scalar multiple of the disease-associated transcriptional dysregulation signature, such that Δ_treat_ = *α* Δ_disease_. An *α* < 0 indicates reversal of the disease effect by treatment, whereas an *α* > 0 indicates exacerbation. The magnitude of *α* for each gene quantifies the extent of reversal and was used to classify genes into five categories: super-reversal (*α* < −1.3); full reversal (−1.3 ≤ *α* < −0.7); partial reversal (−0.7 ≤ *α* < −0.3); negligible reversal (−0.3 ≤ *α* ≤ 0.3); and exacerbation (*α* > 0.3). The transcriptional dysregulation signature was defined in one of two ways: (i) differentially expressed genes (FDR < 0.1) at each corresponding time point in Q111 (empty vector) versus WT (empty vector), or (ii) differentially expressed genes (FDR < 0.1) that were included in a curated panel of 266 striatal genes (STR266) previously reported to be consistently dysregulated across multiple HD mouse models ^38^. We also calculated “reversal correlation of Z statistics” where a positive correlation indicates that *Msh3* KO shifts Q111 transcriptional profiles toward WT-like expression, and “overall reversal fraction,” both as described in Wang et al. ^26^.

To identify gene co-expression patterns, we applied the Weighted Gene Co-expression Network Analysis (WGCNA) framework ^58^. Variance-stabilized normalized counts were used as input. Network topology was evaluated using the pickSoftThreshold function, which calculates the scale-free topology fit index and mean connectivity across a range of candidate powers. A soft-thresholding power of 12 was selected based on the scale-free topology criterion (R² ≥ 0.90). We used a pre-defined set of striatal co-expression modules described by Langfelder et all, including CAG repeat-length–dependent modules derived from an allelic series of Huntington’s disease knock-in mice (Q20, Q80, Q92, Q111, Q140, Q175) ^36^, providing a reference framework for interpreting disease-relevant transcriptional programs. Module eigengenes were correlated with genotype or treatment contrasts to assess the impact of Q111 and *Msh3* KO on network organization. To investigate the potential rescue effect of *Msh3* KO, module-genotype correlations were computed for Q111 (empty) versus WT (empty) and for Q111 *Msh3* KO versus Q111 (empty). Modules exhibiting opposite directions of correlation between these two contrasts were classified as showing reversal (rescue), whereas modules with concordant directions were considered exacerbated.

### *Msh3* knockout effect on *Htt1a* transcripts

To assess the effect of *Ms*h3 KO on *Htt1a* in Q111 mice, HTSeq ^55^ was used to quantify reads aligning to the HD KI hybrid human-mouse intron 1 region ^54^, extending to the previously described cryptic polyadenylation site ^39^ (See Figure 7A). These counts were compared to background intronic signal from the WT allele, estimated by realigning reads from WT animals to the mm39 reference genome and quantifying reads mapping to the corresponding *Htt* intron 1 region up to the cryptic polyadenylation site. Statistical analyses of *Htt1a* expression were performed using 2-tailed unpaired t-tests, comparing to the relevant empty vector-treated groups using TPM-normalized gene expression values.

### Statistical analyses

Statistical analyses of the RNA-seq data are described above. In all other cases, statistical significance was determined by comparing the mean of each *Msh3* KO group with the mean of the respective empty vector group using two tailed unpaired t-tests in GraphPad Prism v10.

## Supplementary Figures

**Figure S1. Relationships between *Msh3* editing, MSH3 protein and striatal expansion**

**A.** *Msh3* editing frequency in striata of Q111 and WT mice determined using a targeted MiSeq assay. Shown are the mean values across 8-10 mice per group. Decrease over age likely in part reflects increased weight **B.** The mean decrease in MSH3 protein (blue; N=3-5 per group) or expansion index (red; N=10 per group) in Q111 *Msh3* guide-treated mice relative to Q111 empty vector-treated mice are plotted against the mean % frameshift editing. **C**. The mean decrease in expansion index in Q111 *Msh3* guide-treated mice relative to Q111 empty vector-treated mice (N=10 per group) is plotted against the mean decrease in MSH3 protein (N=3-5 per group). **D.** Expansion index is plotted against % frameshift editing for individual Q111 mice in the 6-24 and 16-24-week groups (left) and in the 6-40, 16-40 and 24-40-week groups (right). The boxed regions highlight overlapping editing frequencies between treatment conditions (N=10 per group).

**Figure S2. Expansion indices across all groups**

**B.** Quantification of somatic CAG expansion indices in *Msh3* guide-treated, empty vector-treated and untreated mice. Treatment groups are shown below the graphs and analyses ages are shown above. Unt: untreated mice. **** p<0.0001; *** p<0.001 (unpaired Student’s t-tests). N=10 Q111 mice per condition. Bars show mean +/- standard deviation.

**Figure S3. Impact of *Msh3* KO on number of intranuclear inclusions**

**A.** Representative images of striatal sections co-immunostained with anti- HTT mAb5374 (green) and anti-histone H3 (red; merged mAb5374 + histone H3 images are shown), illustrating discrete intranuclear inclusions. Scale bar (white line) = 50 µm. **B**. Quantification of the number of H3-positive nuclei with inclusions (see Methods) in *Msh3* KO and empty vector-treated cohorts analyzed at 40 weeks of age. To control for slide-to-slide variation, the % of nuclei with inclusions in *Msh3 KO* sections was normalized to that in the cohort-matched empty vector-treated section on the same slide, and therefore all empty vector samples have a value of 1 (see Methods). N=4-5 per condition. **** p<0.0001; ** p<0.01 (unpaired Student’s t-tests). Bars show mean +/- standard deviation.

**Figure S4. Impact of the empty vector**

**A.** Numbers of differentially expressed genes (DEGs) (FDR<0.1) elicited by the Q111 allele in mice treated with the empty vector control relative to untreated mice. At each age, the numbers of DEGs are similar, indicating that the vector does not substantially enhance or suppress the response to the Q111 allele. **B**. Numbers of DEGs (FDR<0.1) in WT mice treated with empty vector for each group. **C**. Pathway enrichment of DEGs in WT mice treated with empty vector (16-40 week group is omitted as there were very few DEGs). GeneRatio on the x-axis is the fraction of DEGs represented in each pathway. There are overlaps between the enriched pathways in each group as well as differences in the top enriched pathways, suggesting that the response to the virus may differ depending on the age at which the mouse is exposed to the virus and the time of exposure.

**Figure S5. Reversal correlation and overall reversal fraction**

To assess our results in comparison to other relevant studies we computed “reversal correlation of Z statistics” (left) and “overall reversal fractions” (right) described by Wang et al. ^26^. The reversal correlation of Z statistics uses correlation of gene dysregulation as a measure of transcriptome-wide reversal. *i.e*. it captures the correlation between genes that are dysregulated in one direction in Q111 vs WT and in the opposite direction in Q111 *Msh3* KO vs Q111 (a positive value of this metric means that the genes are altered in the opposite direction). The overall reversal fraction captures the amount of expression normalization using a measure of overall log fold change normalization (a positive value of this metric means that gene expression is normalized to WT). We note that the reversal correlation in the 6-40-week group is between that in *Msh3* null heterozygous and *Msh3* null homozygous Q140 mice, and that the overall reversal fraction in the 6-40-week group is closest to that in *Msh3* null heterozygous Q140 mice^26^. Comparisons of *Msh3* guide-treated Q111 or WT are to empty vector-treated mice.

**Figure S6. Expression levels across *Htt* genotype and *Msh3* KO treatment of full reversal genes.**

The relative effects of *Msh3* KO in the Q111 background and WT background gene expression values (TPM) are plotted for the “full” reversal genes defined by the posterior probability model in the 6-40-week group (A), the 16-40-week group (B) and the 24-40-week group (C). Comparisons of *Msh3* guide-treated Q111 or WT are to empty vector-treated mice.

**Figure S7. Expression levels across *Htt* genotype and *Msh3* KO treatment of partial reversal genes.**

The relative effects of *Msh3* KO in the Q111 background and WT background gene expression values (TPM) are plotted for the "partial" reversal genes defined by the posterior probability model in the 6-40-week group. Comparisons of *Msh3* guide-treated Q111 or WT are to empty vector-treated mice.

**Figure S8. Expression levels across *Htt* genotype and *Msh3* KO treatment of HD modifier and candidate modifier genes**

The relative effects of *Msh3* KO in the Q111 background and WT background gene expression values (TPM) are plotted for the HD modifier and candidate genes in the 6-40-week group. Comparisons of *Msh3* guide-treated Q111 or WT are to empty vector-treated mice.

**Figure S9. Weighted gene co-expression network analysis of non-CAG length-dependent modules**

Heatmap of weighted gene co-expression network analysis (WGCNA) based on striatal non-CAG length-dependent modules defined by Langfelder et al. ^36^ in Q111 vs WT comparisons and in Q111 *Msh3* vs Q111 comparisons for each intervention and analysis age. Red and blue indicate up- and down-regulated modules with p values in parentheses. Comparisons of *Msh3* guide-treated Q111 or WT are to empty vector-treated mice.

**Figure S10. Relationships between nuclear huntingtin intensity, *Htt1a* and mean striatal CAG length.** Top: 24 weeks; bottom, 40 weeks. Left: correlations between anti-HTT mAb5374 nuclear intensity and mean striatal CAG length (left) or *Htt1a* (right) across all groups. N= 4-6 per group. Comparisons of *Msh3* guide-treated Q111 or WT are to empty vector-treated mice.

## DATA AVAILABILITY

Data will be made available upon reasonable request.

## CODE AVAILABILITY

No custom code or software were developed for this study. Details of software packages used are provided in the Methods section.

## ACKNOWLEDGEMENTS

This work was supported by the CHDI Foundation (VCW) and National Institutes of Health (NIH) grant R01 NS049206 (VCW) and R01 NS126420 (RMP). We thank the MGH CCIB DNA Core for assistance with next-generation sequencing; the UMass Chan Medical School Viral Vector Core for AAV production; the Rodent Histopathology Core at Dana-Farber/Harvard Cancer Center (RRID: SCR_009742) for tissue processing, embedding and sectioning (Dana-Farber/Harvard Cancer Center is supported in part by an NIH Cancer Center Support Grant (NIH 5 P30 CA06516)); B. Callahan from the Jackson Laboratory for breeding and genotyping Cas9.Q111 mice; B. Lager at CHDI for mouse logistic support. Figure 1 was generated using BioRender.

## AUTHOR CONTRIBUTIONS

Conceived project: EO, MK, VCW: Devised methodology: EO, MK, ML, AJ, VCW, RMP; Conducted investigations: EO, MK, ML, BJ, FS, NR, JW, AS, RM, TG, EE; Data curation: EO, MK, ML, AJ, VCW; Analyzed the data: EO, MK, ML, AJ, KC, BJ, VCW, RMP; Visualization: EO, MK, ML, AJ, VCW; Project administration: EO, MK, VCW; Supervised the project: EO, MK, VCW; Wrote original draft: EO, MK, AJ, VCW; Manuscript review and editing: ML, KC, BJ, FS, NR, JW, AS, RM, TG, EE, RMP.

