## Supplementary Figures for "Somatic CRISPR editing of *Msh3* mitigates Huntington’s disease pathology in mice"

Figure S1

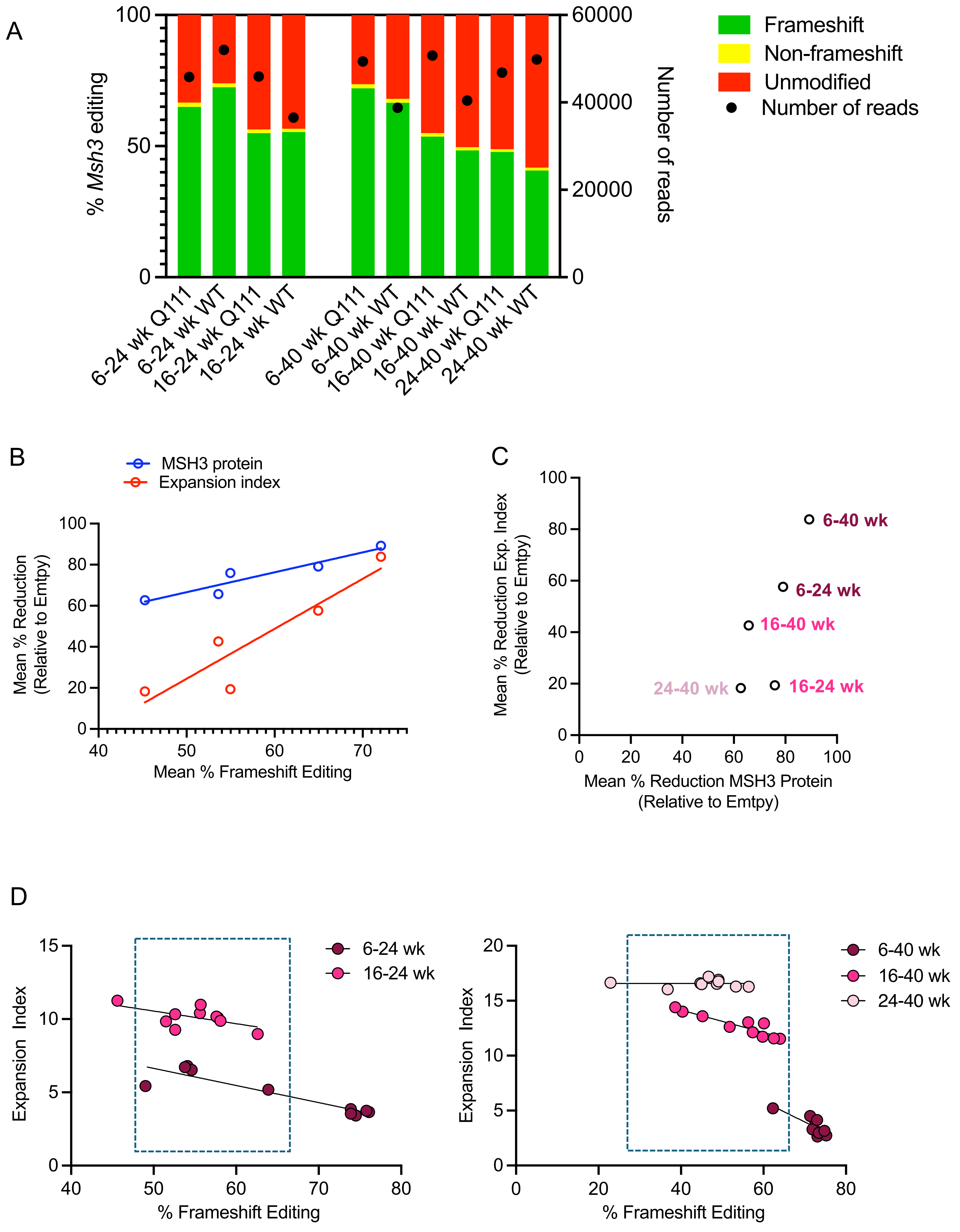

A
% Msh3 editing
Number of reads
Frameshift
Non-frameshift
Unmodified
Number of reads
6-24 wk Q111
6-24 wk WT
16-24 wk Q111
16-24 wk WT
6-40 wk Q111
6-40 wk WT
16-40 wk Q111
16-40 wk WT
24-40 wk Q111
24-40 wk WT
B
MSH3 protein
Expansion index
Mean % Reduction (Relative to Emtpy)
Mean % Frameshift Editing
C
Mean % Reduction Exp. Index (Relative to Emtpy)
Mean % Reduction MSH3 Protein (Relative to Emtpy)
6-40 wk
6-24 wk
16-40 wk
24-40 wk
16-24 wk
D
Expansion Index
% Frameshift Editing
6-24 wk
16-24 wk
6-40 wk
16-40 wk
24-40 wk

Figure S2

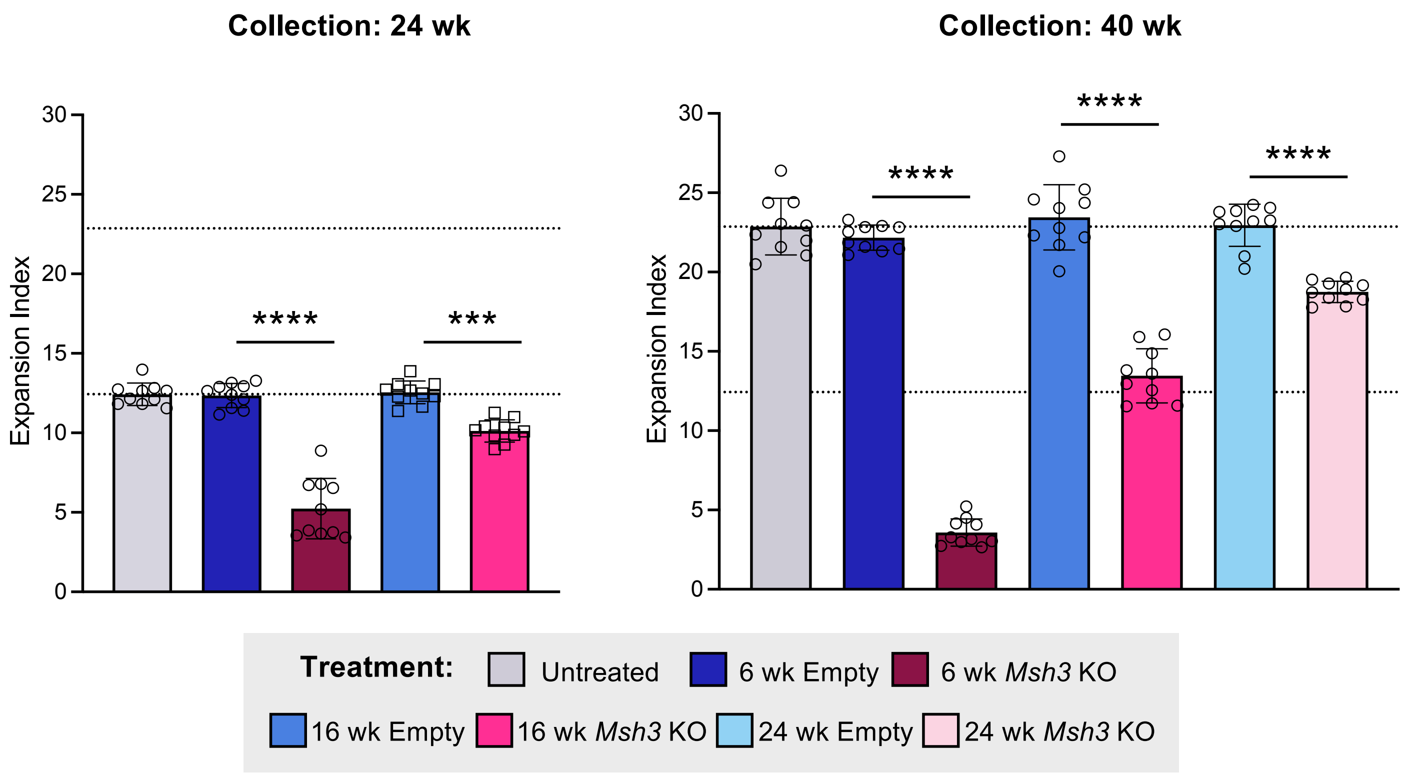

Collection: 24 wk
Collection: 40 wk
Expansion Index
30
25
20
15
10
5
0
****
***
****
****
****
Treatment:
Untreated
6 wk Empty
6 wk Msh3 KO
16 wk Empty
16 wk Msh3 KO
24 wk Empty
24 wk Msh3 KO

Figure S3

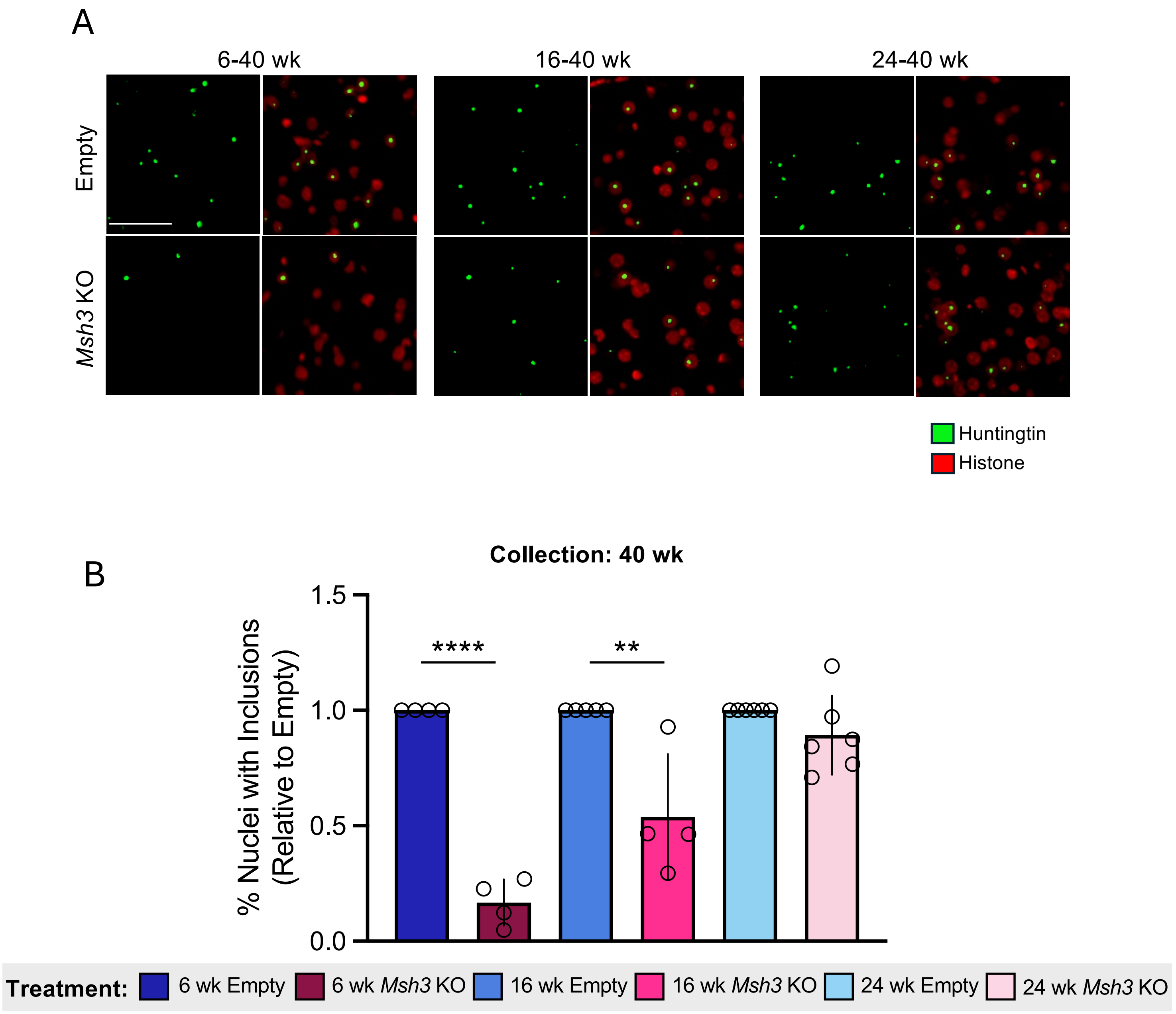
A
6-40 wk
16-40 wk
24-40 wk
Empty
Msh3 KO
Huntingtin
Histone
B
Collection: 40 wk
% Nuclei with Inclusions (Relative to Empty)
1.5
1.0
0.5
0.0
****
**
Treatment:
6 wk Empty
6 wk Msh3 KO
16 wk Empty
16 wk Msh3 KO
24 wk Empty
24 wk Msh3 KO

Figure S4

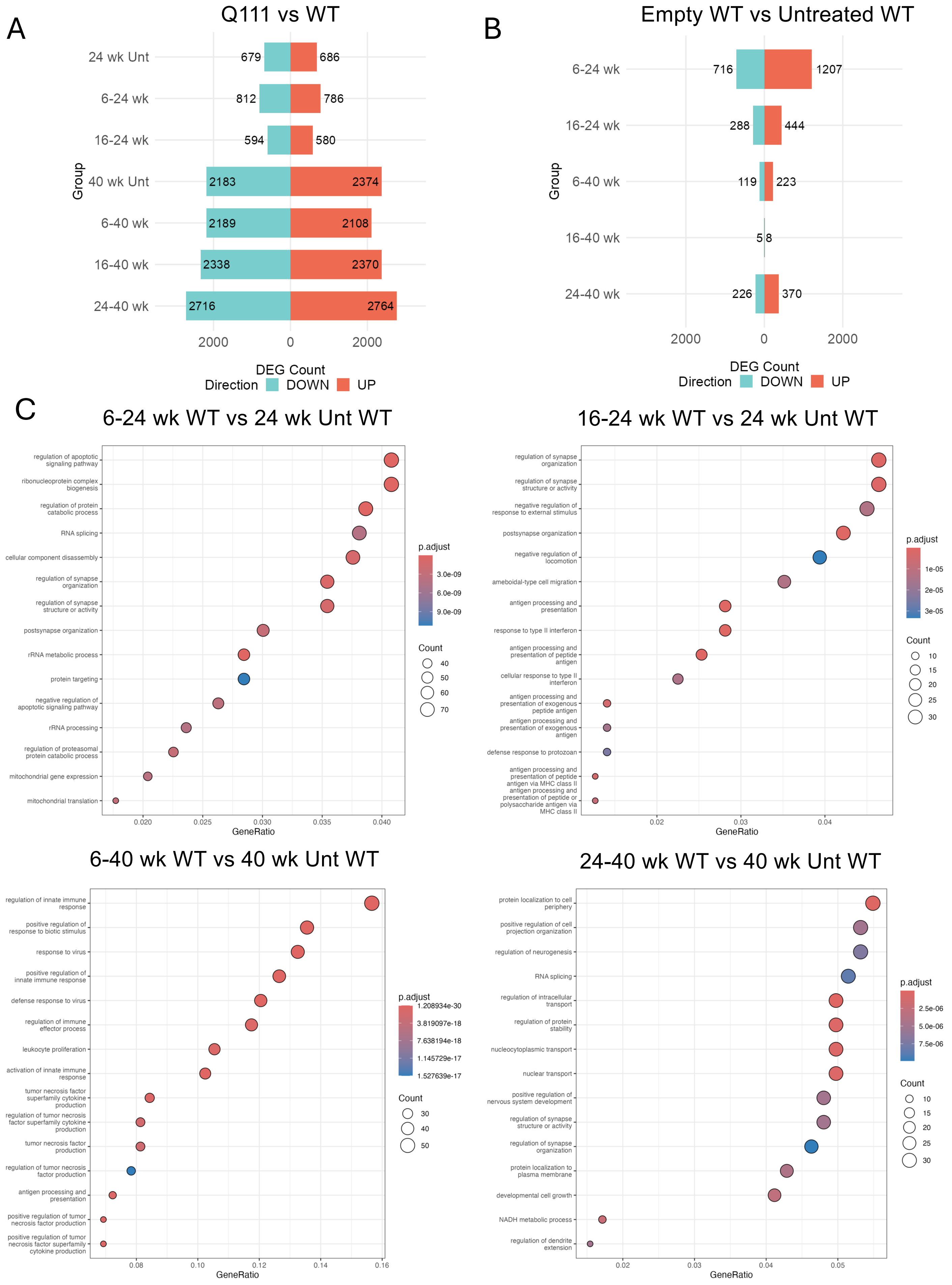
A
Q111 vs WT
24 wk Unt
6-24 wk
16-24 wk
40 wk Unt
6-40 wk
16-40 wk
24-40 wk
679
686
812
786
594
580
2183
2374
2189
2108
2338
2370
2716
2764
Group
2000
0
2000
DEG Count
Direction
DOWN
UP
B
Empty WT vs Untreated WT
6-24 wk
16-24 wk
6-40 wk
16-40 wk
24-40 wk
716
1207
288
444
119
223
5
8
226
370
Group
2000
0
2000
DEG Count
Direction
DOWN
UP
C
6-24 wk WT vs 24 wk Unt WT
regulation of apoptotic signaling pathway
ribonucleoprotein complex biogenesis
regulation of protein catabolic process
RNA splicing
cellular component disassembly
regulation of synapse organization
regulation of synapse structure or activity
postsynapse organization
rRNA metabolic process
protein targeting
negative regulation of apoptotic signaling pathway
rRNA processing
regulation of proteasomal protein catabolic process
mitochondrial gene expression
mitochondrial translation
0.020
0.025
0.030
0.035
0.040
GeneRatio
p.adjust
3.0e-09
6.0e-09
9.0e-09
Count
40
50
60
70
16-24 wk WT vs 24 wk Unt WT
regulation of synapse organization
regulation of synapse structure or activity
negative regulation of response to external stimulus
postsynapse organization
negative regulation of locomotion
ameboidal-type cell migration
antigen processing and presentation
response to type II interferon
antigen processing and presentation of peptide antigen
cellular response to type II interferon
antigen processing and presentation of exogenous peptide antigen
antigen processing and presentation of exogenous antigen
defense response to protozoan
antigen processing and presentation of peptide antigen via MHC class II
antigen processing and presentation of peptide or polysaccharide antigen via MHC class II
0.02
0.03
0.04
GeneRatio
p.adjust
1e-05
2e-05
3e-05
Count
10
15
20
25
30
6-40 wk WT vs 40 wk Unt WT
regulation of innate immune response
positive regulation of response to biotic stimulus
response to virus
positive regulation of innate immune response
defense response to virus
regulation of immune effector process
leukocyte proliferation
activation of innate immune response
tumor necrosis factor superfamily cytokine production
regulation of tumor necrosis factor superfamily cytokine production
tumor necrosis factor production
regulation of tumor necrosis factor production
antigen processing and presentation
positive regulation of tumor necrosis factor production
positive regulation of tumor necrosis factor superfamily cytokine production
0.08
0.10
0.12
0.14
0.16
GeneRatio
p.adjust
1.208934e-30
3.819097e-18
7.638194e-18
1.145729e-17
1.527639e-17
Count
30
40
50
24-40 wk WT vs 40 wk Unt WT
protein localization to cell periphery
positive regulation of cell projection organization
regulation of neurogenesis
RNA splicing
regulation of intracellular transport
regulation of protein stability
nucleocytoplasmic transport
nuclear transport
positive regulation of nervous system development
regulation of synapse structure or activity
regulation of synapse organization
protein localization to plasma membrane
developmental cell growth
NADH metabolic process
regulation of dendrite extension
0.02
0.03
0.04
0.05
GeneRatio
p.adjust
2.5e-06
5.0e-06
7.5e-06
Count
10
15
20
25
30

Figure S5

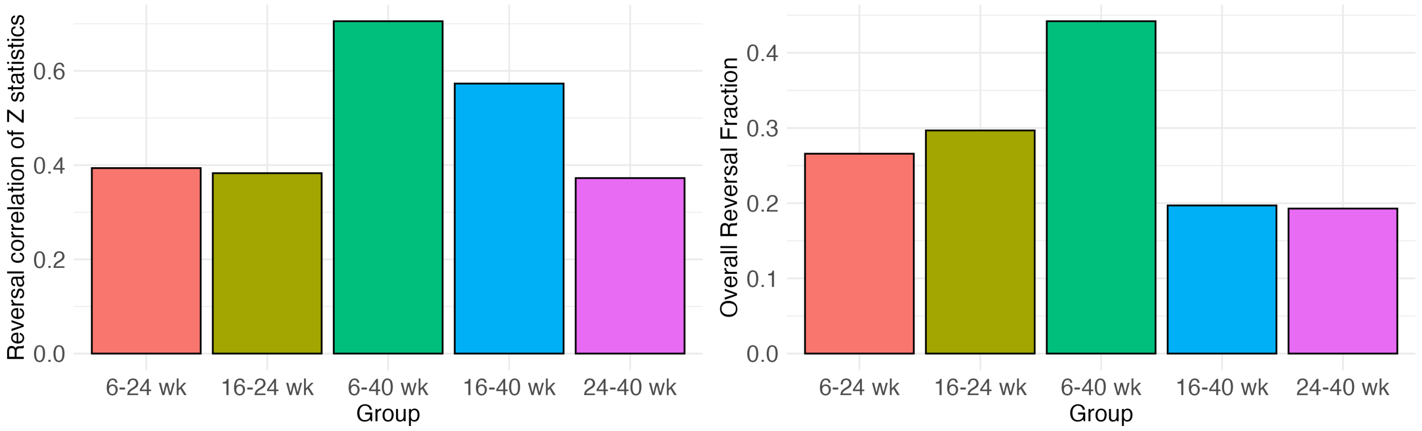
Reversal correlation of Z statistics
0.0
0.2
0.4
0.6
6-24 wk
16-24 wk
6-40 wk
16-40 wk
24-40 wk
Group
Overall Reversal Fraction
0.0
0.1
0.2
0.3
0.4
6-24 wk
16-24 wk
6-40 wk
16-40 wk
24-40 wk
Group

Figure S6

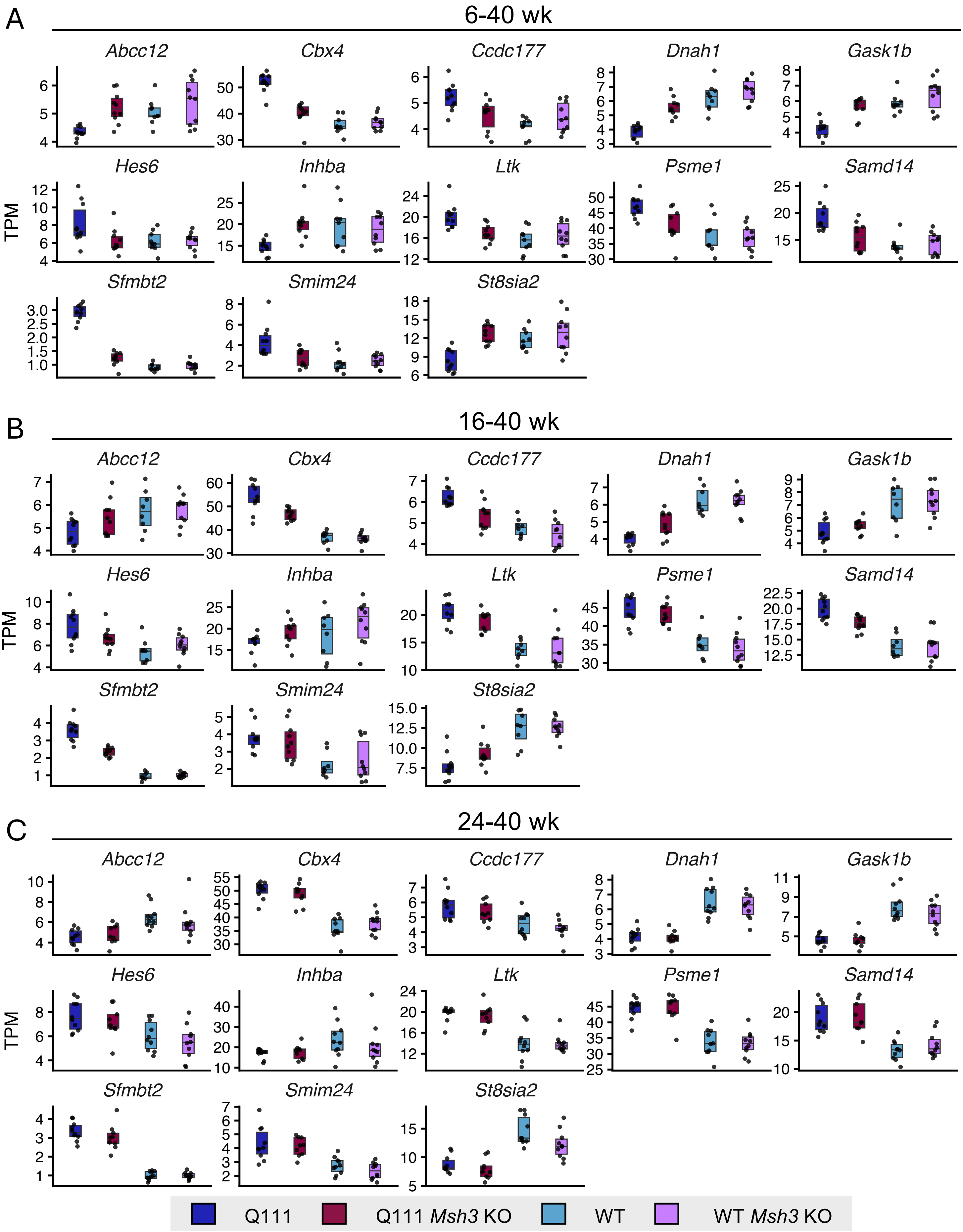
A
6-40 wk
B
16-40 wk
C
24-40 wk
TPM
Abcc12
Cbx4
Ccdc177
Dnah1
Gask1b
Hes6
Inhba
Ltk
Psme1
Samd14
Sfmbt2
Smim24
St8sia2
Q111
Q111 Msh3 KO
WT
WT Msh3 KO

### Figure S7

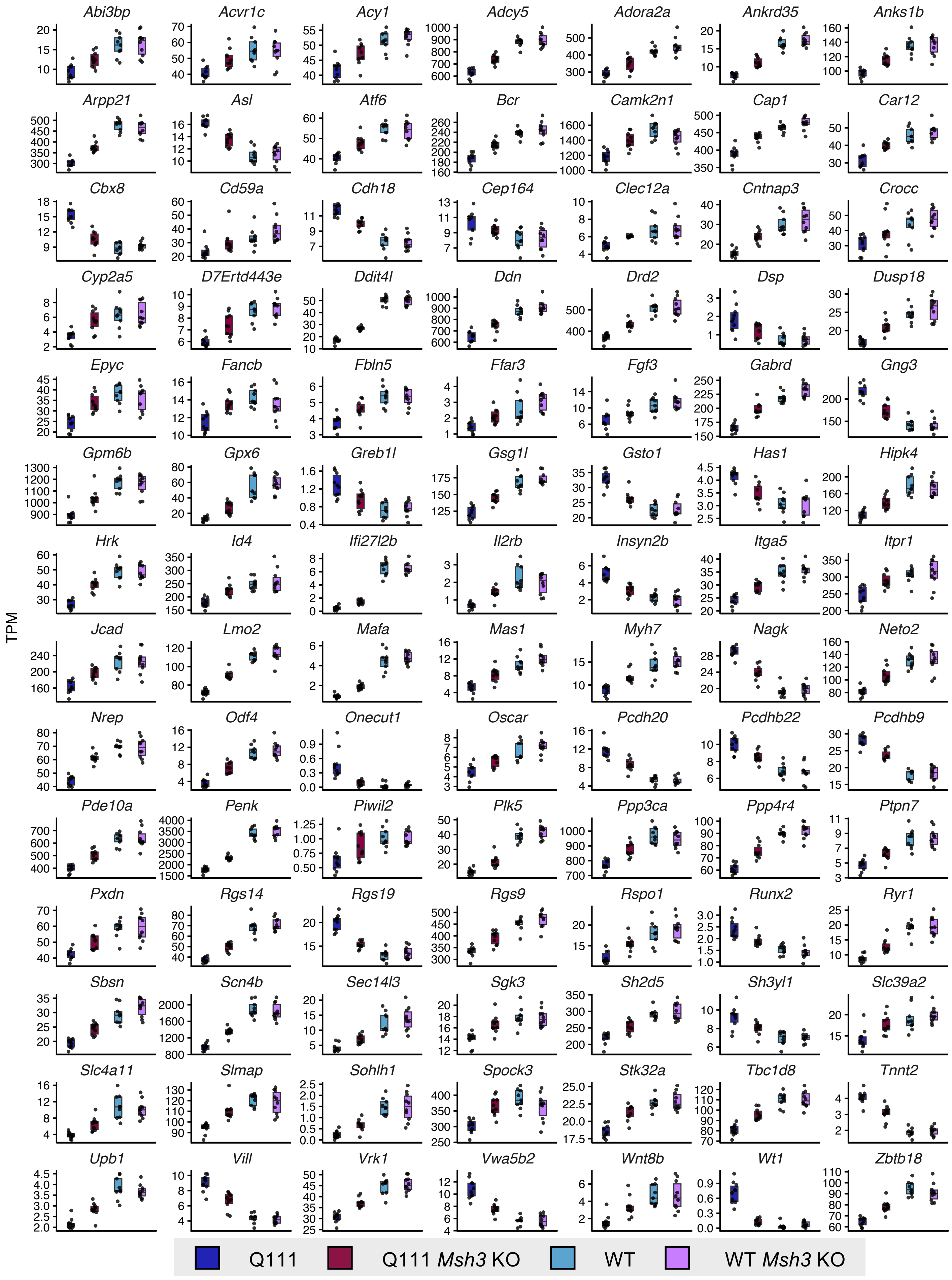
Abi3bp
Acvr1c
Acy1
Adcy5
Adora2a
Ankrd35
Anks1b
Arpp21
Asl
Atf6
Bcr
Camk2n1
Cap1
Car12
Cbx8
Cd59a
Cdh18
Cep164
Clec12a
Cntnap3
Crocc
Cyp2a5
D7Ertd443e
Ddit4l
Ddn
Drd2
Dsp
Dusp18
Epyc
Fancb
Fbln5
Ffar3
Fgf3
Gabrd
Gng3
Gpm6b
Gpx6
Greb1l
Gsg1l
Gsto1
Has1
Hipk4
Hrk
Id4
Ifi27l2b
Il2rb
Insyn2b
Itga5
Itpr1
Jcad
Lmo2
Mafa
Mas1
Myh7
Nagk
Neto2
Nrep
Odf4
Onecut1
Oscar
Pcdh20
Pcdhb22
Pcdhb9
Pde10a
Penk
Piwil2
Plk5
Ppp3ca
Ppp4r4
Ptpn7
Pxdn
Rgs14
Rgs19
Rgs9
Rspo1
Runx2
Ryr1
Sbsn
Scn4b
Sec14l3
Sgk3
Sh2d5
Sh3yl1
Slc39a2
Slc4a11
Slmap
Sohlh1
Spock3
Stk32a
Tbc1d8
Tnnt2
Upb1
Vill
Vrk1
Vwa5b2
Wnt8b
Wt1
Zbtb18
TPM
Q111
Q111 Msh3 KO
WT
WT Msh3 KO

Figure S8

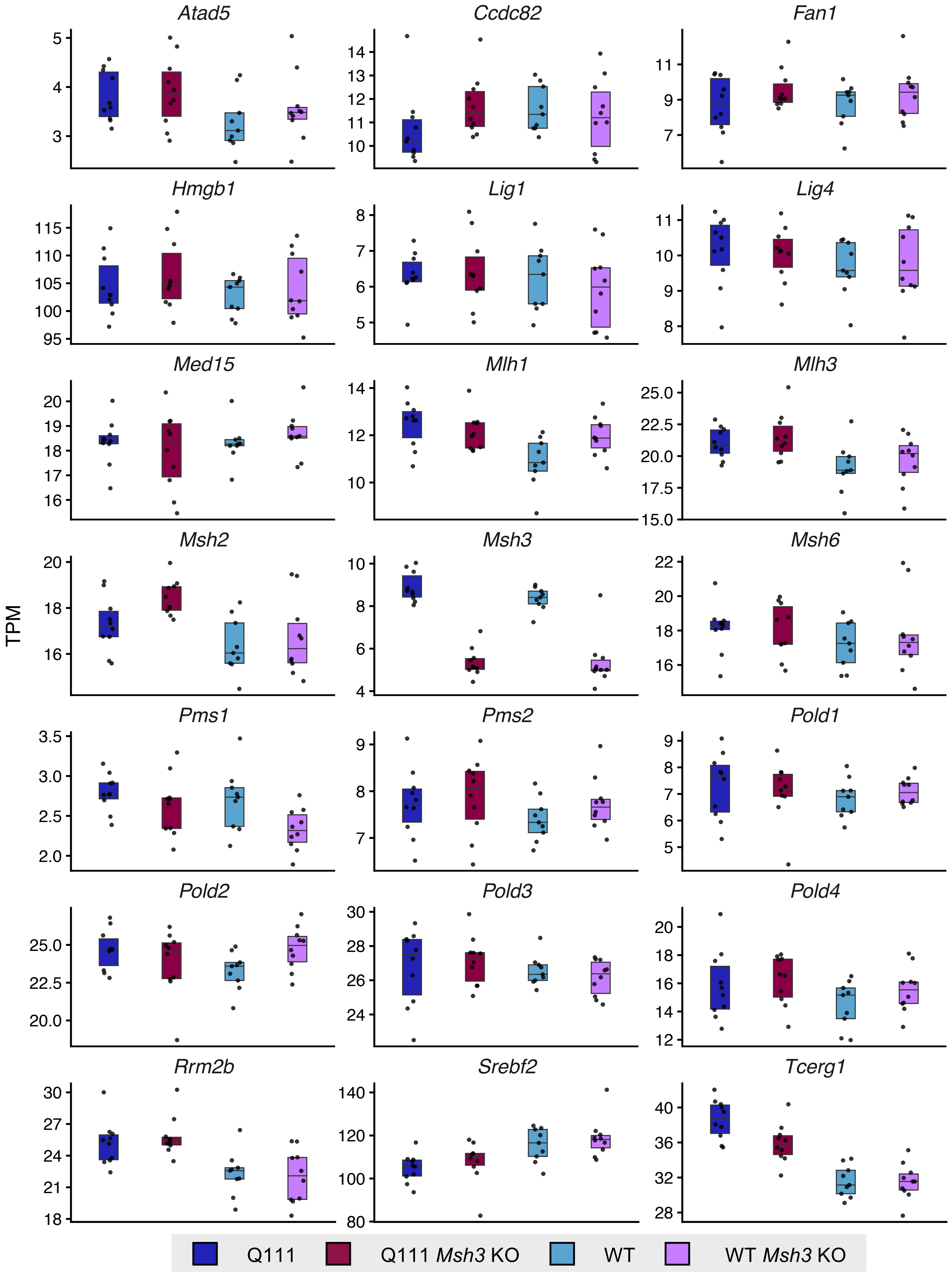

Atad5
Ccdc82
Fan1
Hmgb1
Lig1
Lig4
Med15
Mlh1
Mlh3
Msh2
Msh3
Msh6
Pms1
Pms2
Pold1
Pold2
Pold3
Pold4
Rrm2b
Srebf2
Tcerg1
TPM
Q111
Q111 Msh3 KO
WT
WT Msh3 KO

Figure S9

| | 6-24 wk Q111 vs WT | 6-24 wk Q111 *Msh3* KO vs WT | 16-24 wk Q111 *Msh3* KO vs WT | 6-40 wk Q111 vs WT | 6-40 wk Q111 *Msh3* KO vs WT | 16-40 wk Q111 *Msh3* KO vs WT | 24-40 wk Q111 *Msh3* KO vs WT |
|---|---|---|---|---|---|---|---|
| M4 | -0.051 (0.4) | -0.087 (0.2) | -0.12 (0.07) | -0.085 (0.2) | 0.091 (0.2) | -0.37 (7e-09) | -0.13 (0.05) |
| M5 | -0.1 (0.1) | 0.04 (0.5) | 0.23 (4e-04) | 0.19 (0.004) | -0.26 (6e-05) | -0.15 (0.02) | 0.14 (0.04) |
| M12 | -0.092 (0.2) | 0.14 (0.04) | 0.15 (0.03) | -0.45 (6e-13) | -0.15 (0.02) | 0.12 (0.08) | 0.21 (0.001) |
| M13 | 0.38 (2e-09) | -0.26 (7e-05) | 0.1 (0.1) | 0.18 (0.006) | 0.4 (2e-10) | 0.061 (0.4) | -0.082 (0.2) |
| M14 | -0.43 (6e-12) | 0.21 (0.001) | -0.31 (2e-06) | -0.24 (3e-04) | -0.16 (0.01) | 0.091 (0.2) | -0.051 (0.4) |
| M16 | -0.021 (0.8) | -0.062 (0.3) | 0.032 (0.6) | -0.35 (6e-08) | 0.45 (8e-13) | 0.26 (7e-05) | -0.0053 (0.9) |
| M19 | -0.064 (0.3) | -0.16 (0.01) | -0.32 (6e-07) | 0.3 (4e-06) | -0.13 (0.04) | -0.21 (0.002) | -0.19 (0.004) |
| M22 | 0.032 (0.6) | -0.0045 (0.9) | 0.2 (0.002) | 0.39 (7e-10) | -0.49 (2e-15) | 0.026 (0.7) | 0.15 (0.03) |
| M23 | -0.035 (0.6) | 0.073 (0.3) | 0.12 (0.07) | 0.17 (0.01) | -0.37 (9e-09) | 0.056 (0.4) | 0.096 (0.1) |
| M24 | 0.17 (0.008) | 0.026 (0.7) | 0.098 (0.1) | 0.26 (6e-05) | 0.066 (0.3) | -0.22 (0.001) | -0.061 (0.4) |
| M30 | -0.085 (0.2) | -0.07 (0.3) | -0.21 (0.002) | -0.061 (0.4) | 0.2 (0.002) | -0.066 (0.3) | -0.34 (9e-08) |
| M38 | -0.014 (0.8) | -0.00013 (1) | -0.25 (1e-04) | 0.35 (4e-08) | -0.16 (0.02) | -0.064 (0.3) | -0.29 (9e-06) |
| M40 | -0.1 (0.1) | 0.09 (0.2) | 0.17 (0.008) | -0.24 (2e-04) | -0.15 (0.02) | 0.12 (0.06) | 0.16 (0.01) |
| M41 | 0.34 (1e-07) | -0.089 (0.2) | -0.12 (0.07) | 0.21 (0.002) | 0.3 (3e-06) | 0.14 (0.03) | -0.16 (0.01) |
| M42 | -0.35 (4e-08) | -0.019 (0.8) | -0.37 (6e-09) | -0.0096 (0.9) | -0.088 (0.2) | -0.16 (0.02) | -0.0092 (0.9) |
| M44 | 0.14 (0.03) | 0.11 (0.09) | 0.39 (1e-09) | -0.18 (0.007) | 0.31 (1e-06) | 0.075 (0.3) | 0.085 (0.2) |
| M45 | 0.36 (2e-08) | -0.066 (0.3) | 0.18 (0.005) | 0.43 (8e-12) | -0.36 (3e-08) | -0.028 (0.7) | 0.061 (0.4) |
| M47 | -0.16 (0.02) | 0.097 (0.1) | 0.16 (0.01) | -0.48 (6e-15) | 0.24 (3e-04) | 0.15 (0.02) | 0.14 (0.04) |
| M48 | -0.14 (0.04) | -0.018 (0.8) | -0.11 (0.09) | -0.34 (2e-07) | 0.41 (1e-10) | 0.043 (0.5) | -0.14 (0.04) |
| M49 | -0.32 (5e-07) | 0.038 (0.6) | 0.081 (0.2) | -0.49 (2e-15) | 0.049 (0.5) | -0.41 (1e-10) | 0.25 (1e-04) |
| M50 | -0.043 (0.5) | 0.0092 (0.9) | 0.25 (1e-04) | -0.22 (9e-04) | -0.067 (0.3) | 0.1 (0.1) | 0.28 (2e-05) |
| M51 | -0.11 (0.1) | 0.025 (0.7) | -0.18 (0.007) | -0.38 (2e-09) | 0.27 (3e-05) | 0.055 (0.4) | -0.23 (4e-04) |
| M53 | -0.00045 (1) | 0.12 (0.07) | 0.36 (3e-08) | -0.5 (6e-16) | 0.41 (6e-11) | 0.31 (2e-06) | 0.13 (0.05) |
| M54 | -0.044 (0.5) | 0.023 (0.7) | 0.18 (0.007) | 0.17 (0.009) | -0.33 (2e-07) | -0.18 (0.005) | 0.24 (2e-04) |

Figure S10

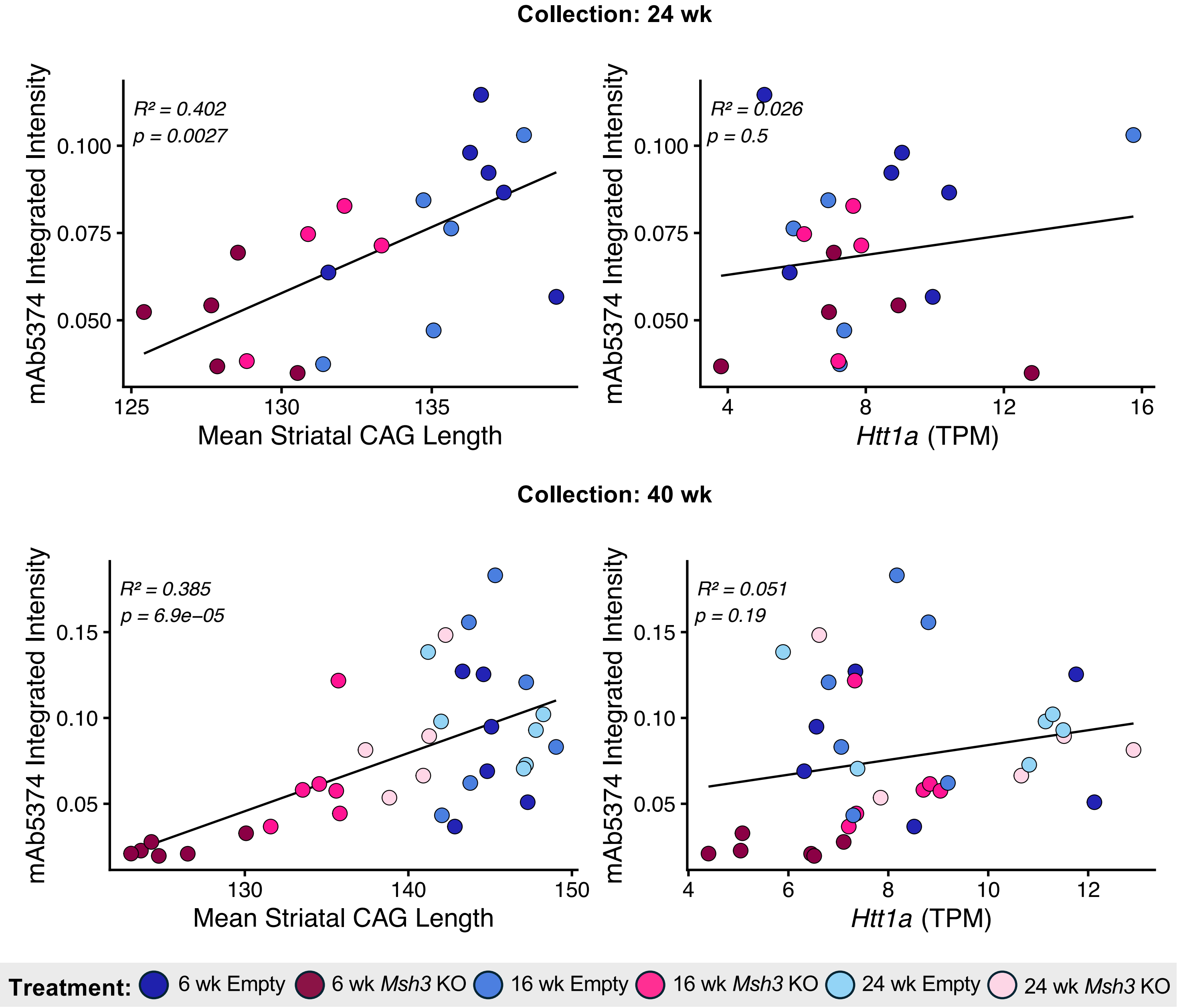
Collection: 24 wk
mAb5374 Integrated Intensity
R² = 0.402
p = 0.0027
0.100
0.075
0.050
125
130
135
Mean Striatal CAG Length
R² = 0.026
p = 0.5
4
8
12
16
Htt1a (TPM)
Collection: 40 wk
R² = 0.385
p = 6.9e−05
0.15
0.10
0.05
130
140
150
Mean Striatal CAG Length
R² = 0.051
p = 0.19
4
6
8
10
12
Htt1a (TPM)
Treatment:
6 wk Empty
6 wk Msh3 KO
16 wk Empty
16 wk Msh3 KO
24 wk Empty
24 wk Msh3 KO
